# Capturing early human memory consolidation utilizing the higher functional specificity of 7T compared to 3T-fMRI

**DOI:** 10.1101/2022.02.01.478615

**Authors:** Silke Kreitz, Angelika Mennecke, Laura Konerth, Julie Rösch, Armin M. Nagel, Frederik B. Laun, Michael Uder, Arnd Dörfler, Andreas Hess

**Affiliations:** Department of Neuroradiology, University Hospital Erlangen, Friedrich-Alexander-University Erlangen-Nürnberg (FAU), Erlangen, Germany; Institute for Pharmacology and Toxicology, Friedrich-Alexander-University Erlangen-Nürnberg (FAU), Erlangen, Germany; Institute of Radiology, University Hospital Erlangen, Friedrich-Alexander University Erlangen-Nürnberg (FAU), Erlangen, Germany

## Abstract

Functional magnetic resonance imaging (fMRI) visualizes brain structures at increasingly higher resolution and better signal-to-noise ratio (SNR) as field strength increases. Yet, mapping the BOLD response to distinct neuronal processes continues to be challenging. Here, we performed 3T and 7T-fMRI analysis of motor-task activation and resting-state connectivity with adjusted SNR. We then applied graph theory to analyze resting-state neuronal networks detected by fMRI after a simple motor task. Despite adjusted SNR, 7T achieved a higher functional specificity of the BOLD response than 3T-fMRI. Following the motor task, 7T-fMRI therefore enabled the detection of an ‘offline replay’ that was directly linked to brain regions associated with memory consolidation. These findings reveal how memory processing is initiated even after simple motor tasks and begins earlier than previously shown. Thus, the superior capability of 7T-fMRI to detect subtle functional dynamics promises to improve diagnostics and therapeutic assessment of neurological diseases.

## Introduction

Magnetic-resonance-imaging (MRI), the gold standard in medical research and clinical diagnostics, generates high resolution images that help diagnose brain injuries, such as aneurysms, stroke and traumatic brain injury, but also tumors and nervous system disease. With the recent clinical approval of ultra-high field 7 Tesla (7T) magnets in 2017, hopes are arising that this technological advance would significantly improve static MRI diagnostics but also functional MRI (fMRI). Specifically, because fMRI detects a blood-oxygen-level-dependent (BOLD) signal coupled to underlying neuronal activity, it could in principle be applied towards mapping and diagnosing human brain function in a range of nervous system indications. In clinical context, monitoring resting-state inter-regional neural activity correlations (rs-fMRI) has several advantages over task-fMRI (mapping static neural activity following a specific motor or sensory task): rs-fMRI data acquisition is less complex, less time consuming and does not need patient’s cooperation. However, the implication of rs-fMRI in clinical practice is currently limited to pre-surgical planning, largely due to the increased complexity required for mapping functional connectivity and analyzing single subjects^1^.

In principle, a stronger magnetic field that improves upon signal-to-noise ratio (SNR) and contrast-to-noise ratio (CNR), enables visualization of finer structures. Accordingly, BOLD 7T-fMRI allows detection of increased BOLD contrast with lesser physiological responses due to a shortened T2* relaxation (the decay of transverse magnetization; an important image contrast determinant)^2,3^, with higher spatial specificity (the intra-vascular signal contribution from draining veins is reduced)^4,5^. However, 7T-fMRI also has disadvantages such as increased inhomogeneity in the static (B0) magnetic field, which can cause visual artifacts that lead to degradation of image quality in a variety of applications.

Since the motor cortex is less influenced by imaging-associated technical artifacts, given its superficial location and proximity to the fMRI coil, motor tasks are frequently used to investigate the field strength influence on neuronal activation in healthy subjects^6,7^ and tumor patients^8^. Indeed, at higher field strength, increased BOLD signal, higher amplitudes and average t-scores as well as increased activated volume have been reported. However, whether a 7T-fMRI analysis of the resting-state connectivity in and of itself, would have potential diagnostic capability, remains largely unexplored. Earlier studies reported that 7T, but not 3T, by significantly higher temporal SNR ratio, improves upon spatial specificity in connected areas^8^. This improvement enabled detection of functional connectivity (FC) that differed in the ventral tegmental area of patients with depression relative to healthy controls^9,10^. However, assessing whether a motor task directly impacts on subsequent resting-state connectivity is beyond the detection limit of 3T-fMRI, and have not yet been demonstrated for 7T-fMRI.

The “replay” of task-related neuronal activity, detected by fMRI in response to auditory and olfactory cues, is thought to contribute to the reconfiguration of memory-related functional connectivity across the brain^11,12^. For 3T-fMRI, this replay was demonstrated during slow-wave-sleep in humans via cue-induced replay of event-related brain activation after declarative memory-encoding tasks^13,14^. Occurrence and strength, especially of hippocampal replay patterns, were correlated with subsequent memory performance^15,16^. Similarly, two human 3T-fMRI studies investigated whether FC-modulation in the resting-state was associated with a previous declarative learning task^17,18^. Although these studies detected a few modulated connections in predefined regions activated during encoding, these were not correlated with memory performance and not controlled for the natural variation of the participant’s resting-state between consecutive measurements. In contrast, for a procedural memory task such as sequential finger-tapping, 3T-fMRI did not detect a replay pattern during sleep^14^ or in the resting-state. Of note, sleep improved the movement speed of a motor task, but not its accuracy^14,19^. Furthermore, analysis of the resting-state revealed enhanced connectivity in executive and cerebellar networks after motor learning but not after motor movement in and of itself^20,21^. Therefore, whether it is possible to detect a replay response coupled to a preceding motor task remains unknown.

Through the use of invasive electrophysiological methods, the existence of ongoing neuronal firing immediately after a motor task has been demonstrated in animals^19,22,23^. This so-called neuronal “offline replay” is presumed to be directly linked to memory encoding, as suppression of the offline replay results in poorer memory performance^23^. Recently, learning-related offline replay in the human brain was reported in a pilot trial in which participants were implanted with intracortical microelectrode arrays^24^, providing proof of principle that early memory encoding in the resting-state can be detected.

In this study, we set out to investigate general characteristics of 7T-fMRI in the human brain, but we also sought to examine whether 7T offers a higher BOLD specificity for examining motor task-driven neuronal events impacting on the connectivity in the subsequent resting-state. To make use of the higher specificity of 7T-fMRI, we specifically sought to detect a functional correlate of the neuronal offline replay in the resting-state that would represent a connectivity modulation after a finger-tapping motor task.

## Results

Eighteen healthy right-handed participants (19-55 yrs, 8 female and 10 male) underwent fMRI measurements at both 7T and 3T. Each fMRI session started and ended with resting-state measurements. During sessions, participants either remained at rest or executed an active right-hand motor task. The motor task was a sequential finger-tapping in a classical block design (14 sec tapping and 14 sec rest, repeated 7 times, totaling 3.5 min). Therefore, the 7T and 3T sessions, and the start and end resting-states (rs1 and rs2) were paired, the rest-group and the active task group (ft-group) were unpaired (Extended Data Fig. 1).

### 7T and 3T functional data can be differentiated mainly by spatial quality metrics

To evaluate the measurement quality for each session, we assessed rs1 across all participants for spatial (average brain volume over time) and temporal (voxel-wise time-courses) quality metrics. To evaluate the spatial data quality, we determined the SNR, the CNR between grey and white matter, the foreground to background energy ratio (FBER) and the entropy focus criterion (EFC). Conversely, we assessed temporal data quality using the temporal SNR (tSNR), the standardized per-image standard deviation of the temporal derivative (zDVARS), the median distance index (MDI), and finally the global correlation (Gcorr). For details see Methods.

Overall, those metrics generated high quality fingerprints (Fig. 1a) that, with the exception of Gcorr, differed significantly between 7T and 3T (Fig. 1a,b, paired t-test, p < 0.05). When we did a principal component (PC) analysis using all metrics, we found an excellent separation of field strengths along the first PC (Fig. 1c) with highest absolute loadings for CNR and EFC in the spatial dimension. In contrast, temporal metrics did not contribute before the third PC and did not account for field-strength dependent group separation (Fig. 1c, Supplementary Table 1), which means that, under the constraint of adjusted resolutions to compensate for field strength-dependent SNRs, temporal data quality is less influenced by higher field strength than spatial image quality. We found that whole brain average SNR and tSNR were even higher for 3T. As expected, the highest tSNR of the 3T measurements was located within the cortex, which was closer to the f-MRI head coil. In contrast, the tSNR of 7T measurements was instead more evenly distributed throughout the brain (Extended Data Fig. 2). Taken together the higher cortical tSNR at 3T did not survive the correction for multiple comparison, whereas subcortical regions and the cerebellum showed a significantly higher tSNR at 7T (Fig. 1d, paired t-test, corrected, p<0.05).

**Fig. 1:**
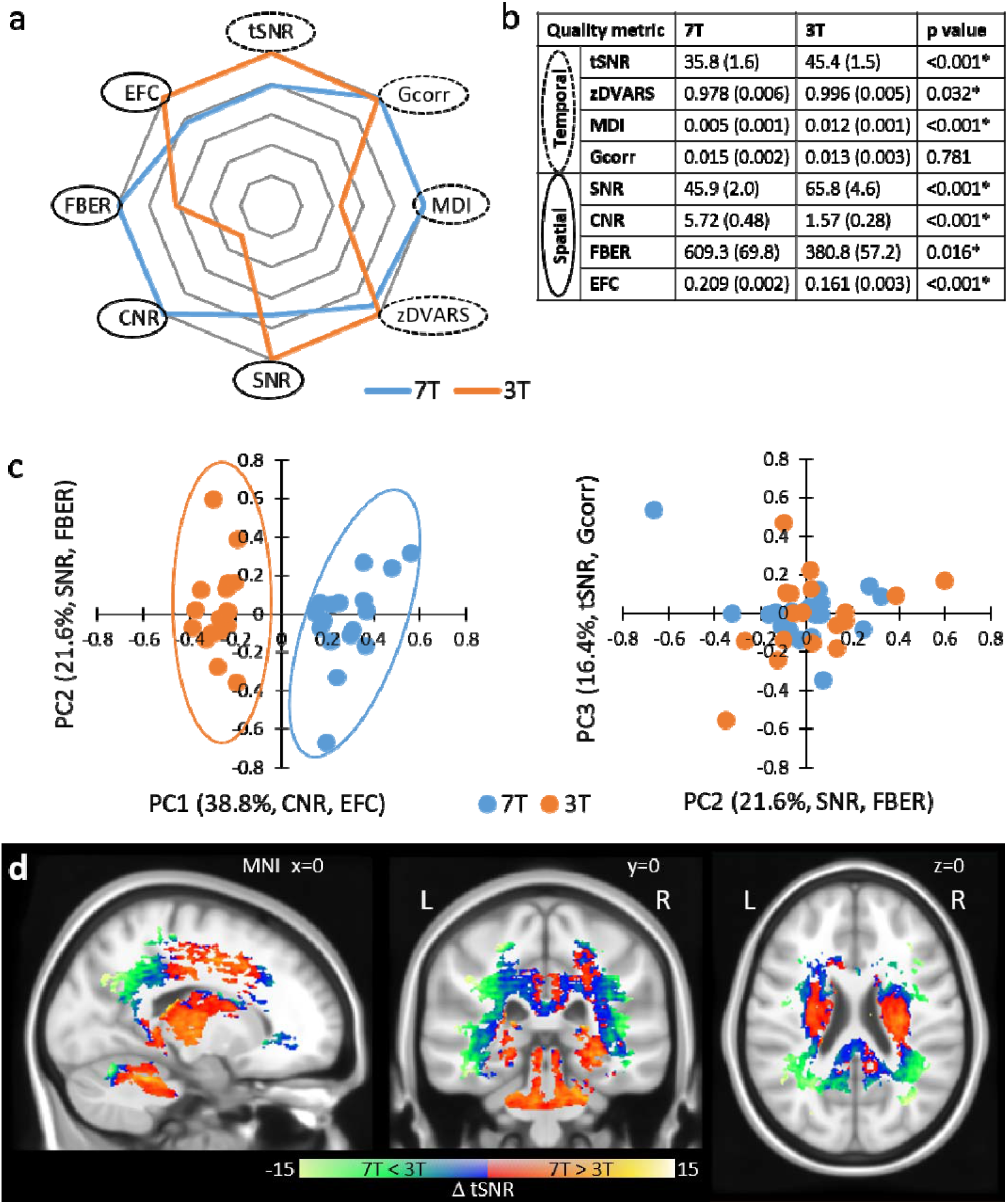
Quality metrics of the first resting-state scan rs1. a) Fingerprints of average quality metrics for 7T and 3T (solid line spatial, dashed line temporal metric). The better value of 7T or 3T is set to 1.0 and the corresponding one is given proportionally. b) Mean quality measures with standard deviation in brackets (n=18). Significance was determined by paired two tailed t-test (*p<0.05). c) Principle Component Analysis (PCA) of quality metrics. Projections of single subject measurements on the first and second PC (left) and the second and third PC (right). Percent eigenvalue and metrics with absolute loadings above 0.5 on the respective PC are given in brackets of axis titles. For detailed loadings of the PCs, see Supplementary Table 1. d) Spatial distribution of significant differences in tSNR between 7T and 3T (n=18, two tailed paired t-test with permutation correction, p<0.05).

### 7T-fMRI revealed a higher motor task related specificity of the BOLD response

Next, we evaluated how field strength influences a BOLD response stimulated by a motor task (n=9, paired design). As an indicator for the region’s activation specificity, we expressed the motor task-induced activation per brain region as percentage of participants, the so-called activation probability (Extended Data Fig. 3). In further analysis, we only included regions in which the activation probability was 100% in at least one hemisphere and with one field strength. The resulting 43 brain regions (21 bilateral and 1 middle) and their corresponding activation probabilities are shown in Supplementary Table 2. Within these regions, the activation probability, i.e. their specificity, was significantly higher with 7T compared to 3T (two factor ANOVA without replication, F=2.441, p=0.002).

We found highly similar spatial activation patterns for 7T and 3T, especially for contralateral sensorimotor areas (Fig. 2a), but the overall activated volume was slightly higher for 7T compared to 3T (F=4.857, p=0.028, repeated measure ANOVA). The average BOLD signal response profile (Supplementary Fig. 1) revealed significantly higher (F=334.955, p=1.6×10^−59^) and broader (F=7.048, p=0.008) response amplitudes with a tendency towards a longer decreasing phase (higher amplitude symmetry), but no difference in time to peak after stimulation onset (Fig. 2b, Supplementary Table 2). Additionally, we calculated the temporal contrast-to-noise ratio (tCNR) per region and also found a significant main effect for field strength (7T > 3T, F=489.46, p=0.000). To identify parameters that contribute the most to a field strength-dependent measurement separation, we performed a PCA using all BOLD response parameters and the tCNR, each averaged per subject over all brain regions. Subject’s measurements were separated according to field strength along the first and the second PC. The first two PCs explained 65 % of the variability between all subjects and loadings above 0.5 were tCNR on the first, and amplitude symmetry and time to peak on the second PC (Fig. 2c). However, we found that the activated volume did not contribute to the data variability or separation between measurements according to field strength (Supplementary Table 3). When we investigated region-specific effects of field strength on brain regions using ANOVA, we did not find any interactions except for amplitude height (F=2.009, p=0.0004) and tCNR (F=3.081, p=1.47×10^−8^). Here, the contralateral primary motor (M1 left) and somatosensory (S1 left) regions showed significantly enhanced response amplitudes (Tukey HSD, corrected, p<0.05,) indicating the high task-related specificity of the BOLD response (Fig. 2d top). The tCNR was enhanced in the left S1 and M1 regions (Fig. 2d bottom), and additionally in bilateral secondary motor cortex and the deep nuclei of the cerebellum (data not shown, Tukey HSD, corrected, p<0.05).

**Fig. 2:**
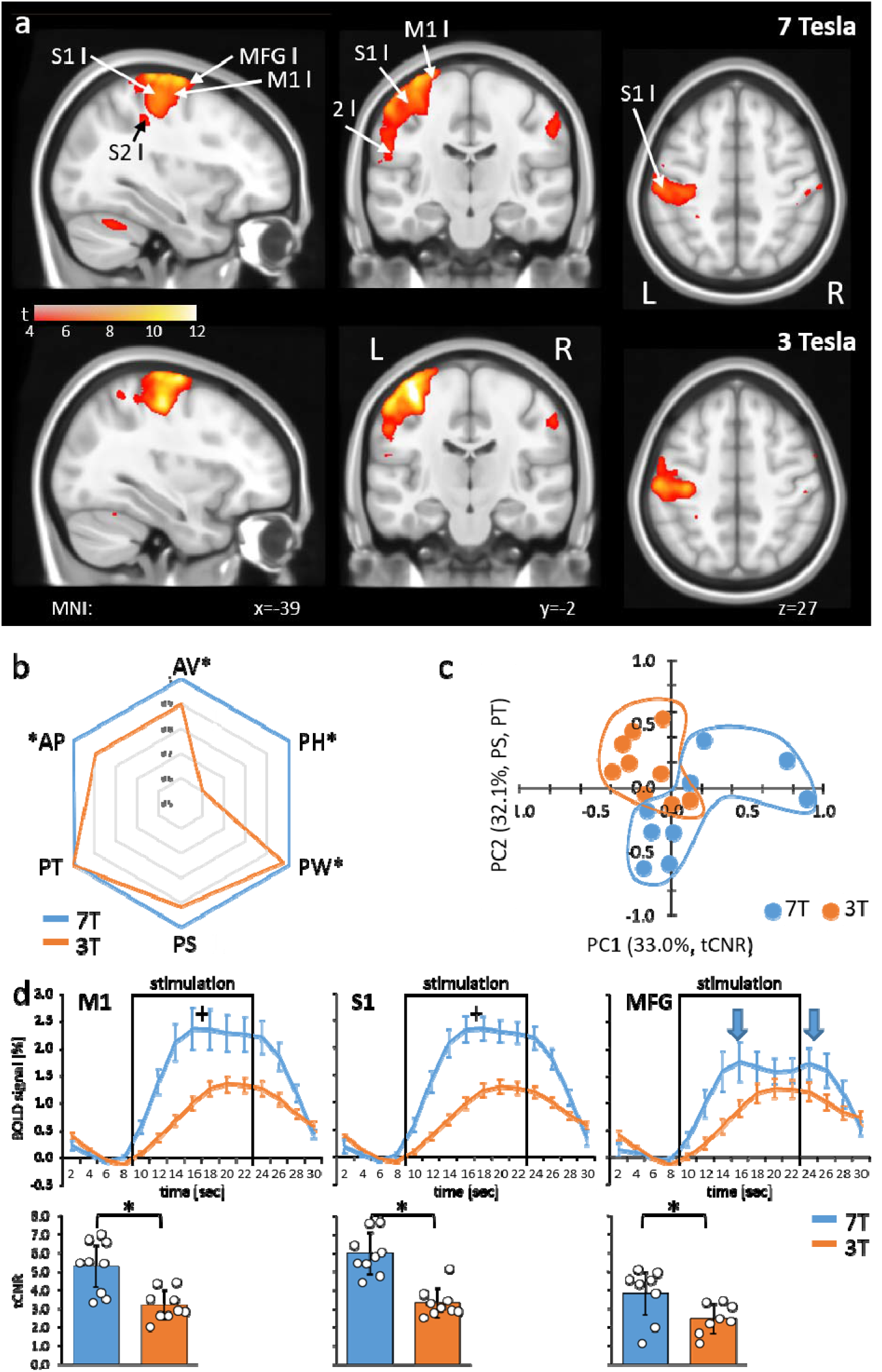
BOLD response to a finger-tapping motor task. a) Neurological view of average statistical parametric maps, thresholded for significant activation using false discovery rate according to Benjamini & Yekutieli (BY-FDR, q=0.05, n=748 time points), acquired for each subject with 7T (top) and 3T (bottom), respectively (n=9). b) Fingerprint plot (left) of normalized average BOLD response parameters (n=9). Significance between 7T and 3T was determined by mixed repeated measure ANOVA without replication for AP and with replication for all other parameters using field strength as within factor and brain regions as between factor (*p<0.05). c) PCA of average BOLD response parameters per subject (n=9). Projections of single subjects on the first and second PC are shown. Percent eigenvalue and the BOLD response parameters with absolute loading above 0.5 on the respective PC are given in brackets in axis titles. For detailed loadings of the PCs, see Supplementary Table 3. d) Average BOLD response profiles (top) of left primary motor cortex (M1), primary somatosensory cortex (S1) and middle frontal gyrus (MFG) and the corresponding temporal contrast-to-noise ratios (tCNR, bottom). Squares mark the stimulation time period. Arrows indicate biphasic amplitude shape. ^+^p<0.05, Tukey HSD with Bonferroni correction, *p<0.05, two tailed paired t-test with Bonferroni correction. AP: activation probability, AV: activated volume, PH: amplitude (peak) height, PS: amplitude symmetry, PT: time to peak, PW: amplitude width (see Supplementary Fig. 1).

### 7T resting-state networks showed higher specificity explicitly in cognitive and sensorimotor networks

To analyze resting-state data, we applied GIFT, a MATLAB tool for performing independent component analysis (ICA) on fMRI data, on the rs1 period (n=18, paired design) for both field strengths. Specifically, we checked for common resting-state networks (RSN) and differences in spatial distribution and network level functional connectivity strength between 7T and 3T. To identify RSNs, we compared the similarity of 20 ICA aggregate components derived from the GIFT group ICA to two sets of publicly available templates^25,26^ (Supplementary results, Extended Data Fig. 4).

In all, we detected nine RSNs as follows: the default mode (DMN), anterior Saliency (aSN), sensorimotor (SMN), left and right executive control (LECN and RECN, respectively), auditory (AuN) and three visual networks: medial or primary (pVN), lateral (lVN) and occipital (oVN) (Fig. 3a). When we cross-correlated the corresponding ICA components to assess the similarity of the RSNs between 7T and 3T, we found that the similarity was highest for the DMN (r=0.85) and lowest for the SMN (r=0.54) with a median value for all nine RSNs of r=0.611 (Fig. 3b).

**Fig. 3:**
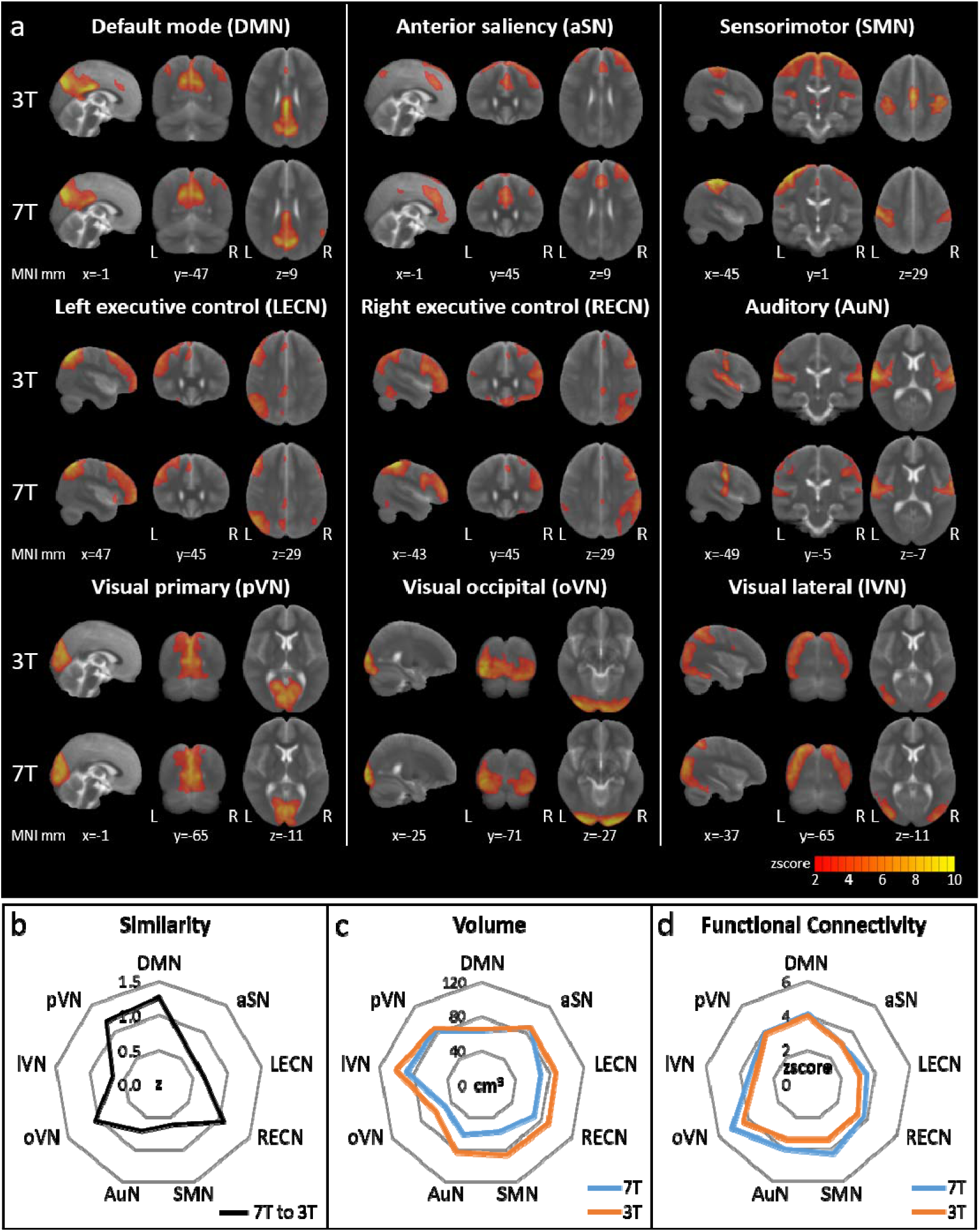
Resting-state Networks (RSNs) derived from 3T and 7T fMRI and 20-component group ICA. RSNs were identified in comparison to two published templates (cf. Supplementary results and Extended Data Fig. 4). a) ICA zscore maps thresholded at zscore=2 (corresponding to p<0.05, uncorrected). Visualization of the three most informative orthogonal slices for each 3T/7T pair superimposed on the MNI standard space template image. b) Similarity of 3T and 7T paired RSNs calculated via spatial cross correlation of ICA zscore maps. c-d) Comparison of thresholded ICA maps (zscore > 2) of 3T/7T pairs using total volume (c) and average zscore as a measure for functional connectivity (d).

Next, we determined the global spatial extension of the thresholded RSN aggregate components (zscore > 2, corresponding to p<0.05, uncorrected) and the average zscore as an indicator for network level FC strength. We found that spatial extension and FC were identical at 7T and 3T for DMN, pVN, and aSN (Fig. 3c,d). Those networks, especially the DMN, are considered task free networks associated with the basic function of the resting brain. However, the more cognitive executive control (LECN, RECN) and higher visual (lVN, oVN) networks and the SMN showed smaller spatial extension and enhanced average FC at 7T (Fig. 3c,d), indicating higher spatial specificity of these RSNs at 7T.

### 7T resting-state graphs showed higher specificity of the underlying functional connectivity

Compared to the ICA-derived RSNs, graph-theoretical FC analysis generates deeper insights into the brain-wide information flow. For this purpose, we used the multi-seed-region approach (MSRA^27^) to create subject specific correlation matrices for rs1 using 158 brain regions (n=18, paired design).

We assumed that a strong homotopic connectivity in healthy subjects would be largely independent of its anatomical distance^28^ (see Supplementary results) and did not observe any differences in data quality between 7T and 3T (Extended Data Fig. 5). The specificity^29^ was determined by the ratio of the FC strength between central regions of task negative DMN (anterior cingulate (Cga) and middle cingulate cortex) and between the Cga and the middle frontal gyrus (MFG; a core region of the task positive ECN). With 7T, we found significantly higher specificity compared to 3T (Fig. 4a, paired t-test, p<0.05).

**Fig. 4:**
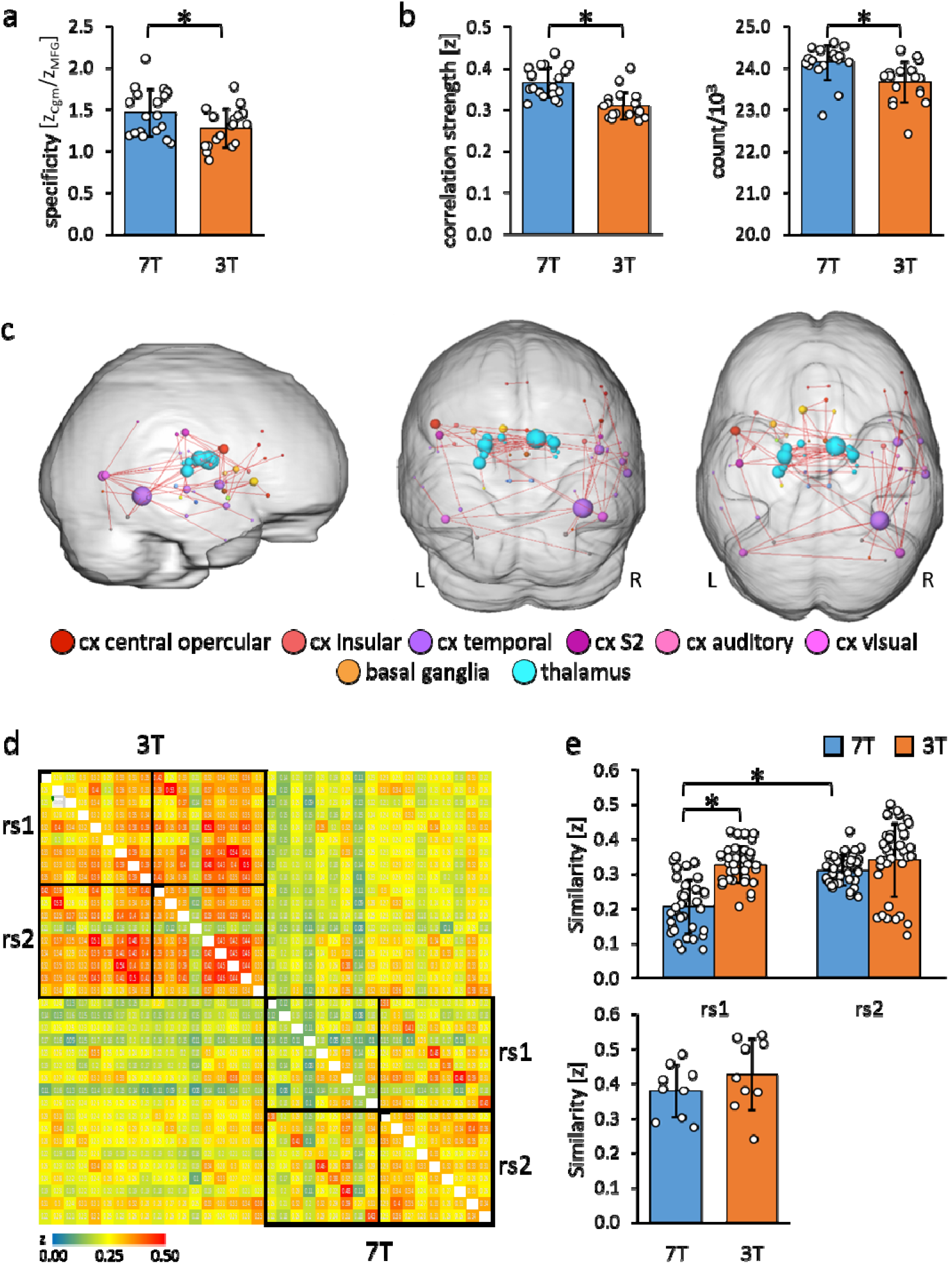
Evaluation of resting-state graphs derived from multi-seed-region analysis (MSRA). a) Specificity of resting-state graphs of the first scan (rs1) described as ratio of the functional connectivity strength (z) of the anterior cingulum to the middle cingulum (Cgm) and to the middle frontal gyrus (MFG) (n=18). Significance was determined by two tailed paired t-test (*p<0.05). b) Properties of rs1 graphs thresholded for significantly correlating connections via BY-FDR (q=0.05, n=298 time points). Left: average functional connectivity strength. Right: number of significant functional connections (n=18). Significance was determined by two tailed paired t-test (*p<0.05). c) Significantly enhanced functional connectivity of 7T compared to 3T rs1 graphs with a connection density of 7% (n=18). Only connections are shown that remain significantly strengthened after each measurement’s normalization to its average functional connectivity strength. Significant components were determined by paired controlled network based statistics (pNBS, α=0.014, p_FWE_<0.0001). d-e) Reproducibility and variability of both resting-state scans rs1 and rs2 at 3T and 7T. Only participants with rest in between both scans were considered (n=9). d) Similarity matrix of all resting-state graphs assessed via spatial cross correlation of the MSRA z-correlation matrices. e) Subject variability derived from pairwise correlation values. Top: between subject variability (n=36 subject pairs, one factor repeated measure ANOVA followed by Tukey HSD with Bonferroni correction, *p<0.05). Bottom: within subject variability (n=9 rs1/rs2 pairs, two tailed paired t-test, no significance).

### 7T resting-state graph topology showed stronger connections within subcortical brain regions

In general, as determined by FDR, 7T rs1 data revealed more significantly correlating connections and higher average connectivity strength relative to 3T (Fig. 4b, paired t-test, p<0.05). For topological comparisons, the 7% strongest correlations of the average correlation matrices were represented as network graphs consisting of nodes (marking brain regions) and edges (marking functional connections between them). Those graphs can be fractionated into non-overlapping distinct communities (Extended Data Fig. 6). These communities mostly resemble the ICA RSNs, but at a more detailed level, thereby demonstrating the high correspondence of graphs and ICA components (Extended Data Fig. 7).

Next, we compared edge-specific FC strength using paired controlled network based-statistics against an independent control group^27^ (11 healthy subjects that underwent two 3T resting-state scans state scans on different days, with no significant differences observed between scans at the α-level of p<0.014). Thus, statistical differences beyond this p-value are most likely attributed to field strength differences and not by the variations of repeated measures per se. Based on the family wise error (p_FWE_) of the whole component of interconnected nodes, we assessed significant differences beyond this α-level by permutation of pairs between control and study group. To adjust for the general higher correlation values at 7T we normalized the data to the same mean in order to capture specific FC strength enhancements related to rather qualitative effects. As shown in Fig. 4c, we demonstrate that the enhanced connectivity at 7T was maintained for subcortical and inferior regions, particularly for the thalamus, basal ganglia and the temporal cortex (α=0.014, p_FWE_ < 0.0001).

### At 7T, the second resting-state measurements were more harmonized between subjects

Next, we demonstrated the reliability of the resting-state measurements by cross-correlating single subject’s matrices for rs1 and rs2 without the motor task in between (n=9), which were similar for each subject (Fig. 4d). We detected significantly higher variability between subjects at rs1 for 7T compared to 3T (one factor repeated measure ANOVA, F=31.63, p<0.05), but no significant field strength-dependent differences at rs2, likely reflecting a higher specificity required to characterize individual resting-state networks. Additionally, 7T rs2 measurements were more similar between subjects than the corresponding rs1 measurements (Fig. 4e top, ANOVA follow up Tukey HSD, corrected p<0.05,) indicating that the shared environmental conditions (the rest inside the scanner), led to more harmonized resting-states in 7T rs2. This effect was not observed with 3T. The intra-subject reproducibility (n=9) was not significantly different between field strengths (Fig. 4e bottom, paired t-test, p<0.05).

### 7T-fMRI revealed significant modulations in the resting-state after an executed motor task

Given our findings suggesting that 7T-fMRI detects functional signals in cortical regions with a higher tSNR-independent specificity, we next assessed short-term resting-state modulations immediately following a finger-tapping motor task. Subjects who performed the motor task between the two resting-state measurements, each at 7T and 3T, (ft-group, n=9) were compared to those remaining at rest during the whole session (rest-group, n=9). Give the harmonization of the inter-subject variability we detected in rs2, we hypothesized that motor task effects might only be visible in the rs2 group comparison. Indeed, we only detected a significant component of reinforced connections in the 7T rs2 group following the motor task (α=0.01, pFWE=0.003, network-based statistic [NBS] analysis, see Methods). Interestingly, the increased connectivity strength occurred primarily in regions relevant to the previous motor performance, namely M1 and MFG, but now ipsilateral to the stimulation side (Fig. 5a). Both regions were activated bilaterally, though the M1 contralateral region was, as expected, more strongly affected (Supplementary Table 2). To investigate relationships between contralateral activation and ipsilateral connectivity modulation, we calculated the interhemispheric functional connectivity between bilateral M1 and MFG during motor performance. We found that the average time courses of both bilateral regions were significantly correlated (critical z-value=0.072, p<0.05, df=744), indicating a strong interhemispheric communication. However, this correlation was significantly weaker for 3T compared to 7T (Extended Data Fig. 8, paired t-test, p<0.05,).

**Fig. 5:**
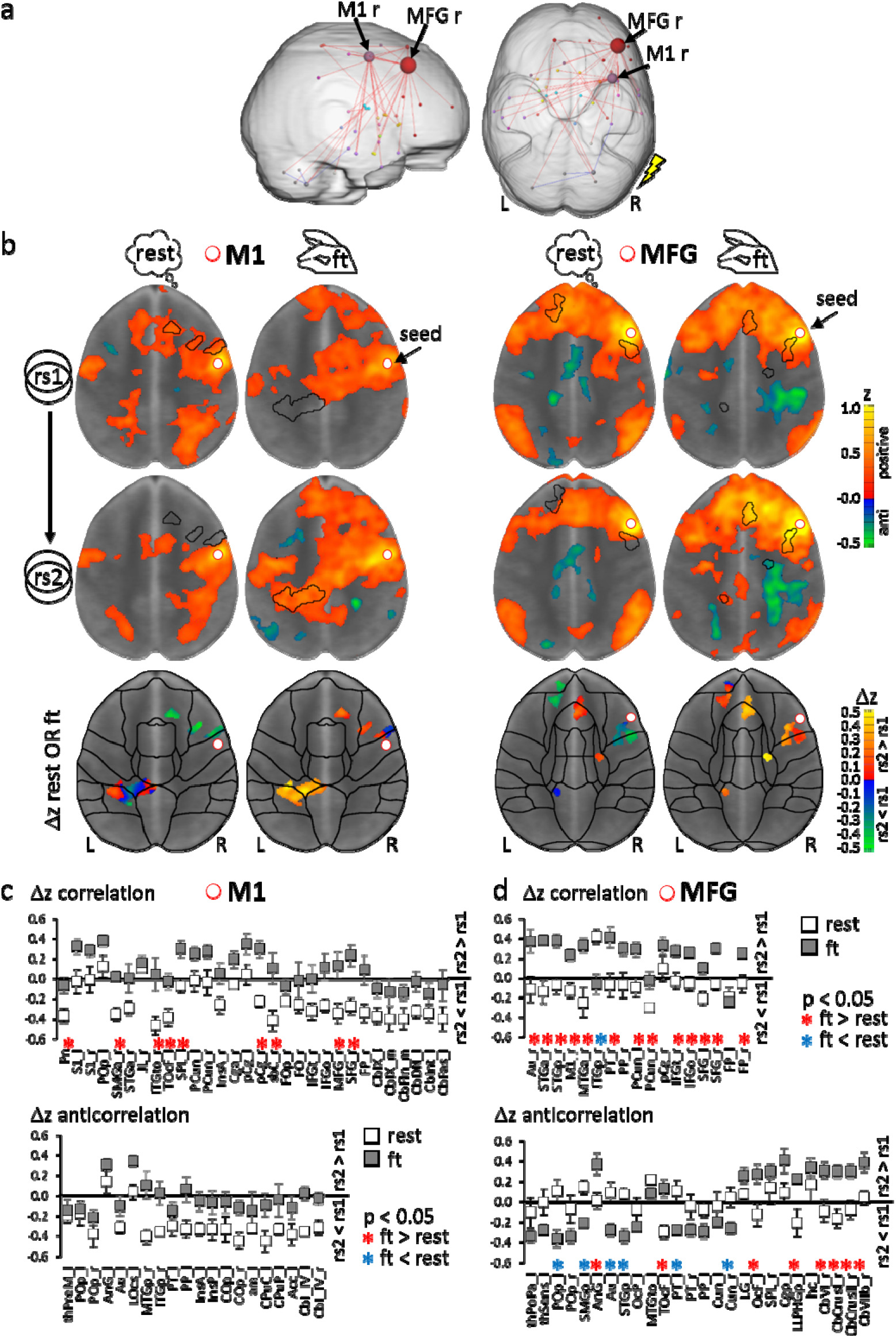
Analysis rational to detect modulations due to a prior motor task performance. a) Significant graph components of enhanced resting-state connectivity strength in the second resting-state scan rs2 after finger-tapping compared to rs2 after rest (n=9 per group, NBS, α=0.01, p_FWE_=0.003). The flash indicates the stimulation side. b) Combined seed region correlation (increasing red to yellow) and anticorrelation (increasing blue to green) maps using seeds in right motor cortex (M1, left) and right middle frontal gyrus (MFG, right) superimposed on MNI standard space template image. Maps were thresholded for significance using BY-FDR (q=0.05, n=298 time points). The presented horizontal slice was chosen with respect to intersect the seed region (MNI z=26 for M1 and z=28 for MFG). Bottom: Areas with significant correlation and anticorrelation differences between rs2 and rs1 in combined for both rest- and ft group. Displayed are average group differences per voxel (two tailed paired voxel wise t-test with permutation correction, p<0.05). c-d) Regional average differences Δz (rs2-rs1) separately for ft- and rest-group (n=9 per group), but based on the same set of voxels derived from b) for seed M1 (c) and seed MFG (d). Only regions with significant voxels exceeding 1% of total region voxels were taken into account. Region-specific significance was determined by two tailed unpaired t-test with Benjamini-Hochberg FDR correction (*p<0.05, red: ft > rest, blue: ft < rest). L: left hemisphere, R: right hemisphere. For abbreviations of region names, see Supplementary Table 2.

Next, we used the M1 and MFG regions at 7T as seeds for further analysis within subjects. For both the right M1 and right MFG, we compared the seed correlation maps of rs1 and rs2 using a voxel wise paired t-test, and identified significant voxels (p<0.05, corrected) separately for the ft- and the rest-group. Subsequently, the significant voxels of both groups were combined and assigned to atlas-based brain regions. Thus, group-specific differences in connectivity strength were assessed within the same set of voxels (Fig. 5b). Finally, using this set of voxels, the average FC differences between subsequent resting-state measurements (Δz rs2-rs1) per brain region in the ft-group were controlled by an unpaired t-test against those in the rest-group (p<0.05, corrected). Higher Δz values in one group indicate a more increased (positive Δz) or less decreased (negative Δz) FC strength in rs2 compared to the other group (Fig. 5c,d).

Ultimately, we found that, controlled against the rest-period, execution of the motor task led to significantly increased Δz of M1 to ipsilateral frontal, secondary somatosensory (S2) areas, contralateral superior parietal cortex, bilateral inferior temporal cortex (ITe) and pons (Fig. 5c top, Fig. 6). We did not observe any significant differences in anticorrelating Δz (rs2-rs1 of negative correlations in the above-described set of voxels, Fig. 5c bottom). In MFG, we found increased Δz to the ipsilateral inferior frontal, auditory, medial temporal (MTe), bilateral superior frontal and precuneus cortex (Fig. 5d top, Fig. 6).

**Fig. 6:**
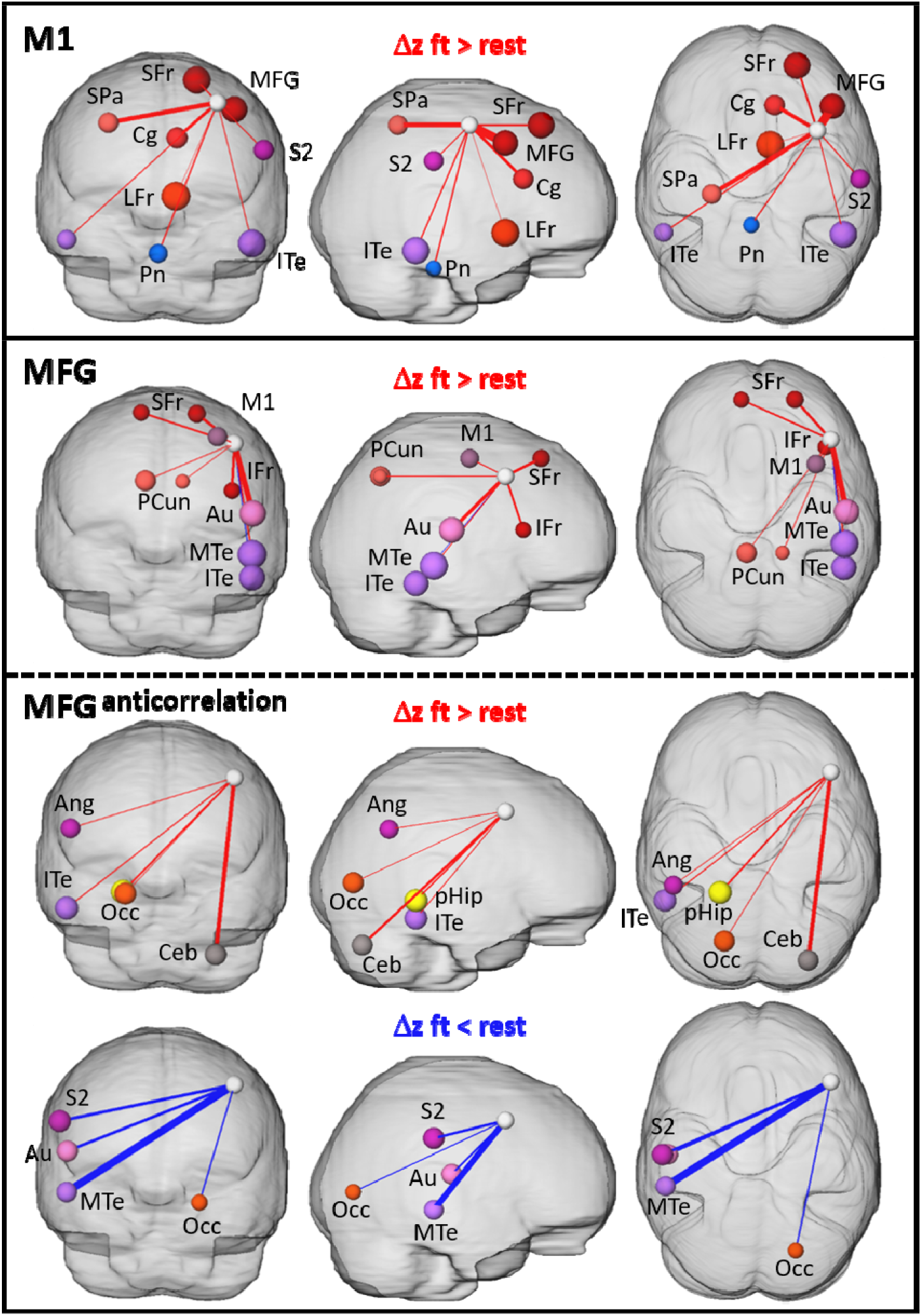
Summary of resting-state connections modulated due to prior finger-tapping. 3D visualization of graphs with nodes and edges. Node size represent absolute difference between Δz (rs2-rs1) of the ft-group and the rest-group, edge thickness indicate the size of the affected area (% voxel of total region voxel). The white node represents the seed (M1 and MFG, respectively). L: left hemisphere, R: right hemisphere. Ang: angular gyrus, Au: auditory cortex, Ceb: Cerebellum superior posterior lobe, Cg: cingulate cortex, IFr: inferior frontal cortex, ITe: inferior temporal cortex, LFr: limbic cortex, M1: primary motor cortex, MFG: middle frontal gyrus, MTe: middle temporal cortex, Occ: occipital cortex, Pcun: precuneus, pHip: parahippocampal gyrus, Pn: Pons, S2: secondary somatosensory cortex, SFr: superior frontal cortex, SPa: superior parietal cortex.

In contrast to the dominantly enhanced positive correlations with ipsilateral regions, anticorrelating Δz involved mainly contralateral regions. Here, the contralateral angular gyrus, ITe, occipital cortex (Occ), parahippocampus and the ipsilateral superior posterior lobe of the cerebellum showed enhanced anticorrelated Δz, and contralateral S2, auditory cortex, MTe and ipsilateral Occ showed reduced anticorrelated Δz in the ft-group compared to the rest-group (Fig. 5d bottom, Fig. 6 bottom). Additionally, both seeds reinforced their connectivity to each other (Fig. 5c,d, Fig. 6).

Taken together, we demonstrate that a unilateral finger-tapping motor task results in contralateral activation of M1 and MFG detected by both 3T and 7T-fMRI. However, modulation of ipsilateral M1 and MFG functional connectivity during the subsequent rest period was only detected by 7T-fMRI.

## Discussion

Here, we examined the influence of high magnetic field strength on fMRI using the BOLD signal as an indirect measure of neuronal processes in the brain. Specifically, our analysis demonstrates that, depending on the field strength, the SNR adapts through a targeted variation of the voxel size. Nevertheless, we successfully demonstrated a higher functional specificity of the BOLD signal with 7T-fMRI.

Motor task-stimulated 7T BOLD responses showed a starkly increased response amplitude and tCNR, a direct consequence of the field strength-dependent boost of the BOLD signal itself, consistent with previous studies^3,6–8^. Additionally, those studies report a 300 to 400 % increased activated volume indicating largely extended activation patterns (i.e. involved brain regions) and higher spatial sensitivity at 7T. However, in our study with adjusted SNRs for both field strengths, we observed only a 10% increase of the activation volume at 7T, and the overall activation pattern was nearly identical. Thus, we conclude that the field strength-dependent higher SNR in earlier work is the main contributor of the higher spatial sensitivity of the BOLD response.

Functional connectivity in the resting-state can be analyzed by focusing on the 3D domain of interacting network patterns as obtained by ICA, or over time using graph-theoretical approaches. Here, we found that spatial patterns in resting-state networks derived from ICA analysis were highly consistent between 3T to 7T. Interestingly, especially the DMN and the pVN did not only display the highest spatial similarity but also nearly identical volumes and zscores, supporting their general relevance in characterizing functional connectivity during rest. The task-associated cognitive and sensorimotor networks were more variable, and expressed smaller volumes but higher zscores at 7T, reflecting individual characteristics of cognitive and sensorimotor resting-state networks^30^.

In contrast, graph-theoretical approaches such as MSRA, which rely on pairwise correlations of time courses, are capable of resolving functional connectivity over time. Even with lower tSNR at 7T, we successfully identified significantly stronger functional connectivity, indicating enhanced functional specificity. This improvement was largely independent of the spatial activity distribution and characterized by tSNR-independent increased BOLD response amplitude, tCNR and functional connectivity strength. These effects might be caused directly by the field strength-dependent enhanced sensibility to the paramagnetic deoxyhemoglobin. Specifically, the ratio of deoxy/oxyhemoglobin reflects the demand of local energy consumption, which is tightly coupled to the underlying neuronal activity. This idea is supported by the strong correlation between local field potential (LFP) and BOLD signal changes. Notably, LFP also reflect energetically expensive synaptic activity^31^. Furthermore, Siero et al. (2014)^32^ demonstrated that the 7T gradient-echo BOLD signal is strongly correlated with the underlying electrophysiology and thus corresponds well with the neuronal activity. Therefore, we conclude that the increased functional specificity at 7T was caused by the closer association of the BOLD signal with the underlying neuronal activation in the brain.

To exploit the increased functional specificity of 7T-fMRI, we investigated resting-state modulations after a motor movement paradigm. Even under the constraint of a completely data-driven analysis, we identified the cortical regions M1 and MFG, activated by the motor performance, as the core regions that were specifically modulated in the subsequent resting-state. Importantly, these modulations could not be detected with 3T, providing proof of concept that 7T detects resting state-specific modulations, directly driven by neuronal activity that is reflected as specific functional connectivity. We propose that these modulations represent a functional correlate to the offline replay of neuronal firing sequences, detected by using invasive electrophysiology, e.g. for the mouse auditory cortex after an auditory task^23^ or the human motor cortex after a motor task^24^. The concept of detecting neuronal replay by fMRI was very recently shown by Wittkuhn et al.^33^ in the human visual cortex using task-related stimuli with various inter-stimulus intervals. Frequency spectra analysis of probabilistic patterns in pre- and post-task resting-states revealed differences at frequencies indicative for the fastest and slowest stimulus presentation speed, pointing towards a replay of activation patterns in the visual cortex^33^.

In this study, we present a neuronal replay visible in brain-wide functional connectivity modulations, complementing the findings of replay of electrophysiological neuronal firing sequences^24^ and probabilistic fMRI activation patterns^33^ in humans. In contrast to those previous studies, we were able to characterize function and nature of the replay. The majority of modulated functional connectivity did not resemble that of brain regions participating in previous finger tapping-associated motor processing, but instead corresponded to frontal-parietal and temporal circuits involved in memory processing and consolidation^34–37^. This was unexpected, since finger-tapping is presumed to predominantly lead to motor area activation not overlapping with working memory regions^38^. Moreover, task-induced neuronal representations of brain activity were found to be inherent in the resting-state^39^, reflecting at least some task-induced brain activity patterns during rest^40,41^. However, the offline replay correlate we identified by 7T-fMRI, indicates that replayed neuronal firing sequences immediately initiate the modulation of functional connectivity related to memory circuits – dominantly working memory such as the frontal-temporal parietal circuit but also regions with central roles in autobiographic (here the precuneus^42^) and episodic (here the medial temporal gyrus^43^) memory.

We also detected modulated anticorrelations in MFG, an important region in memory processing, but not in M1 that dominantly reflects the former motor activation. The switch between task-positive and task-negative (i.e. anticorrelated) resting-state networks is an important feature in memory consolidation^44–47^ (see Supplementary discussion). Additionally, the switch between hemispheres regarding M1 might reflect a hemispheric encoding/retrieval asymmetry^48–51^. Here, M1 was activated on the left hemisphere during finger-tapping (respective encoding) and showed enhanced functional connectivity to memory related regions on the right hemisphere in the subsequent resting-state, which can be presumed to be an early maintenance phase of memory consolidation (see Supplementary Discussion).

In conclusion, we demonstrated, for the first time, the application of 7T-fMRI to detect a highly specific neuronal activity in the resting-state that is directly linked to the execution of a motor task. In combination with our sophisticated analysis workflow, this advance might open the door to more detailed and sophisticated basic research aimed at understanding human memory consolidation but also neurological diagnostics in clinical routine in the very near future.

## Methods

### Participants

A total of 18 (8 female, 10 male) healthy right-handed participants aged 19–55 yrs (average 37 yrs) were recruited. 11 participants had previous experience in being MRI scanned. Exclusion criteria included occurrence of any current or past form neurological/psychiatric diseases or having any contradictions to fMRI scanning. Ethical approval (189-15B) was provided by the local ethics committee of FAU, and informed consent was obtained from all participants. The study adhered to the tenets of the Declaration of Helsinki.

### Study design

To compare the effects of high magnetic fields on functional MRI we conducted a paired study design. Each participant underwent one 3T and one 7T measurement, respectively. The measurement order was balanced between all participants. The time between both measurements ranged from 1 to 6 weeks. Each session consisted of two resting-state measurements (rs1 and a time-offset rs2). In between, half of the participants performed a motor stimulus driven BOLD experiment (ft-group), the other half remained at rest (rest-group, Fig. 1). ft- and rest-group allocations were randomized under the constraint of balanced gender, age, measurement order and previous MR experience.

### fMRI stimulation paradigm

The motor stimulation was a right hand sequential tap of each digit with the thumb lasting 14 sec. This finger-tapping stimulation was repeated 7 times with baseline intervals of 14 sec and additional 14 sec baseline before the first finger-tapping. Thus, the duration of the whole stimulation sequence was 3 min 30 sec. The participants were instructed to start and stop the finger-tapping by voice commands.

### Acquisition

MRI Scans were performed on Siemens Magnetoms TERRA (7T) and TRIO (3T) using 32 channel head coils. Prior to the functional scans an anatomical MP2RAGE scan and a gre-field mapping was performed. The following setting were used for MP2RAGE. 7T: TR = 4500 ms, TE = 2.27 ms, GRAPPA 3, 0.8 mm isotropic resolution, TI = 1/3.2 sec, flip angle = 4°, TA = 9:15 min. 3T: TR = 1.9 sec, TE = 2.52 ms, TI = 900 ms, flip angle = 9°, 1 mm isotropic resolution, GRAPPA 2, TA = 4:26 min. Gre-field mapping was done at 7T using a flash sequence (TR = 4.4 ms, TE = 1.02, 3.06 ms, 3.9 mm isotropic resolution, flip angle = 10 °, Grappa 2, TA = 12 s) and at 3T using the Siemens product sequence (TR = 650 ms, TE = 4.92, 7.38 ms, 2 mm isotropic resolution, flip angle = 60 °, TA = 2:27 min). The anatomical scan was solely for clinical purposes on request of the participants. To acquire functional MRI scans we used gradient echo-planar imaging sequences (GE-EPI) with following settings: GRAPPA (3 with 48 reference lines), TR = 2 sec, TE = 21.0 ms (7T) and 29.6 ms (3T), flip angle = 69° (7T) and 73° (3T), acquisition matrix = 168 × 168 (7T) and 126 × 126 (3T), no. of slices = 84 (7T) and 72 (3T), FOV = 252 mm × 252 mm, resolution = 1.5 mm^3^ (7T) and 2 mm^3^ (3T) isotrop. The resolution was adjusted to achieve comparable signal-to-noise ratios for 7T and 3T: since it can be assumed that noise increases at least linear with B0^52^, the resolution ratio was chosen to match the inverse field strength ratio, i.e. 1.5^3^/2.0^3^ ~ 3/7. Both resting-state measurements acquired 300 volumes each (total time 10 min) and the sequence between both resting-state, either with or without finger-tapping, contained 105 volumes (total time 3 min 30 sec).

### Preprocessing of functional MRI data

After discarding the first two volumes to avoid MR saturation effects, fMRI data were corrected for slice scan time (cubic spline interpolation considering the scan order table) and motion (trilinear detection and sinc interpolation), and were subsequently smoothed spatially (3D Gaussian filter with FWHM 4 mm). The temporal dimension of the stimulus driven BOLD data was smoothed using a GLM-Fourier-Filter with 2 cycles and that of the resting-state data was band pass filtered with a frequency cut off between 0.009 and 0.08 Hz. All preprocessing steps except the band pass filtering of the resting-state data were performed using Brainvoyager QX (Brain Innovation, Maastricht, Netherlands; V2.8.2.2523). Band-pass filtering and all further analysis, if not stated otherwise, was done with MagnAn (Biocom GbR, Uttenreuth, Germany, V2.5), an IDL application (Exelis Visual Information Solutions Inc., a subsidiary of Harris Corporation, Melbourne, FL, USA, V8.5) designed for complex image processing and analysis with emphasis on (functional) MR imaging.

### Individual brain atlas registration and brain region segmentation

To identify anatomical brain regions, we developed a modified Montreal Neurological Institute (MNI) probabilistic brain atlas that was a combination of the Harvard-Oxford-cortical-and-subcortical atlas including white matter and ventricles^53^, the fsl-oxford-thalamic-connectivity atlas^54^ and the SUIT cerebellum atlas^55^. Additionally, a skilled neuroanatomist manually divided the brainstem into medulla, pons, left and right tegmentum, and left and right tectum. In total, this atlas spanned 166 brain regions plus white matter and ventricle regions. We used the first volume of each functional MRI sequence (with the skull stripped manually) as an anatomical reference, which we registered to the linear ICMB152 T2 template in MNI space using a diffeomorphic registration algorithm provided by Advanced Normalization Tools (ANTS^56^, http://stnava.github.io/ANTs/). Subsequently, the resulting individual transformation matrices and fields were applied backwards on each volume of the probabilistic atlas, i.e. each brain region. The median-filtered (kernel 3) maximum probability maps in the individual space of each participant were then used to define brain regions for further analysis. We focused on grey matter brain regions for resting-state data analysis, leaving white matter and ventricles as additional ROIs. Finger-tapping stimulation data were analyzed using brain regions covering both grey and white matter but excluding ventricles.

### Quality metrics

The preprocessed first resting-state scan (rs1) was used to determine the following quality metrics:

Signal-to-noise ratio (SNR)^57^: The mean intensity within the gray matter divided by the standard deviation of the values outside the skull. Higher values are better.
Contrast-to-noise ratio (CNR)^57^: The difference of gray matter and white matter mean intensity values divided by the standard deviation of the values outside the skull. Higher values are better.
Foreground to background energy ratio (FBER)^58^: The variance of intensity values inside the brain divided by the variance of intensity values outside the skull. Higher values are better.
Entropy focus criterion (EFC)^59^: The Shannon entropy of volume voxel intensities proportional to the maximum possible entropy for a same sized volume. This quality metric indicates ghosting and head motion induced blurring. Lower values are better.
Temporal signal-to-noise ratio (tSNR)^60^: Voxel wise calculated mean signal over time divided by the standard deviation over time, yielding tSNR volume maps. Higher values are better.
zDVARS^61^: DVARS is the standard deviation of the temporal derivative of the data, calculated as the spatial standard deviation of the temporal difference image. For better inter-cohort comparisons, DVARS was scaled relative to its temporal standard deviation and autocorrelation. Lower values are better.
Median distance index (MDI)^62^: The mean distance (1-spearman’s rho) between each time point’s volume and the median volume. Lower values are better.
Global correlation (Gcorr)^63^: The average correlation of all pairs of voxel time courses inside the brain indicating global data fluctuations. Values closer to 0 are better.

Grey matter and white matter regions were defined using the corresponding ROIs resulting from the individual brain atlas registration. Background regions were automatically defined as the largest contiguous region of all voxels outside the brain tissue mask with lower intensity than the 5% quantile within the brain mask. Thus, skull and muscles outside the brain tissue mask are reliably eliminated.

### Analysis of finger-tapping stimulation data

Preprocessed finger-tapping stimulation data underwent a classical General Linear Model (GLM) analysis with the hemodynamic response function (HRF) convolved with the boxcar stimulation function as only predictor. This first step was done in Brainvoyager QX, further analysis of the resulting Statistical Parametric Maps (SPMs) was performed in MagnAn. To identify significantly activated voxels, the individual SPMs were thresholded using the Benjamini-Yekutieli version of the False Discovery Rate^64^ (BY-FDR, q=0.05, n=748 time points) which considers a dependency between multiple tests. Since neighboring voxels influence each other this version seems to be the most appropriate to correct for voxel-wise statistical tests. Significantly activated voxels were assigned to distinct brain regions by multiplying the thresholded SPMs with the corresponding maximum probability map. The proportion of participants per group that showed any activated voxels within a certain brain region was defined as the activation probability of that brain region. Under consideration of the different voxel resolutions for 7T and 3T measurements, the number of activated voxels was expressed as activated volumes (mm^3^) per brain region and measurement. For further analysis only those regions were taken into account which exceeded an activation volume larger than 15 mm^3^ and had, under this constraint, an activation probability of 100% either in the 7T or in the 3T group (in total 43 regions, including bilateral counterparts, see Supplementary Table 2). The average time course of activated voxels within these brain regions was extracted and all stimulation periods (“ON” in the boxcar function) including 5 baseline time points (“OFF” in the boxcar function) before and after the stimulation period were averaged. BOLD response was expressed as percent BOLD signal change (ΔR/R) using the average of the three inner time points of the baseline before the stimulation period as baseline reference R. Using this BOLD response amplitude the following parameters were calculated to characterize the stimulation response: Amplitude (peak) height (PH): maximum % BOLD signal change.

Time to peak (PT): time point of maximum % BOLD signal change, expressed in sec after stimulation onset.

Amplitude width (PW): all time points above 75 % of maximum % BOLD signal change, expressed in sec.

Amplitude symmetry (PS): ratio of time points after and before time to peak above whole baseline (before and after stimulation period) mean + standard deviation. Values greater than 1 indicate tailing of the response amplitude, and values lower than 1 mean a slower increase to the maximum. Additionally to those regional parameters we calculated the temporal contrast-to-noise ratio (tCNR^65^) per activated voxel. tCNR was calculated from ΔS_CNR_/σ_t-noise_, with ΔS_CNR_ defined as the following difference:

(mean value of all time points within the stimulation period “ON”) – (mean value of all time points within the baseline period “OFF”).

σ_t-noise_ is the standard deviation of the difference between the original and smoothed signals indicating the non-task-related variability over time. Smoothing was performed using a Savitzky–Golay filter with a polynomial order of 2 and length 5. Voxel-wise tCNR was averaged over activated voxels for each brain region.

### ICA analysis of resting-state data and identification of resting-state networks

Prior to further resting-state analysis the average time courses of white matter and ventricles was regressed out of the preprocessed and band pass filtered resting-state data. White matter and ventricle masks were defined from the individual brain region segmentation as described above.

For ICA analysis, the whole rs1 scans of all participants were registered to the T2 MNI template by repetitively applying the transformation matrices calculated for the individual brain atlas registration. Group ICA analysis of concatenated time series was performed separately for 7T and 3T scans using the FastICA algorithm^66^ in the “Group ICA of fMRI Toolbox” (GIFT v1.3g; https://trendscenter.org/software/). 20 independent components were calculated. For group comparison (7T vs 3T) only the aggregate components were used.

We identified common resting-state networks (RSN) by comparing the 20 GIFT 3T group ICA aggregate components with two different public available template sets provided in the MNI152 space. The first template (https://www.fmrib.ox.ac.uk/datasets/brainmap+rsns/), published by Smith et al. (2009)^26^, is based on images of 36 healthy subjects, who were scanned with a 3T Siemens TRIO Magnetom and analyzed using GIFT group ICA with 20 components. Except for the length of the measurement (6 min instead 10 min in our study), this was the same protocol as we used. Therefore, this template (further on called Smith Template) was most suitable to serve as a reference for the identification of RSNs within our datasets. The second template was provided by the Stanford University (http://findlab.stanford.edu/functional_ROIs.html), and is further on referred to as Stanford Template. The authors^25^ measured 15 subjects for 10 min using a 3T GE Scanner and subsequently calculated a MELODIC group ICA with 30 components. 14 RSNS were identified by visual inspection and were arbitrarily binarized to obtain in total 90 distinct functional ROIs. Two inconsistencies between both templates had to be solved: 1) Due to the higher number of ICA components to create the Stanford template, the DMN was separated into a ventral and a dorsal part. Those two RSNs were combined to one. 2) The Smith Template contained three networks, named executive control and left and right frontoparietal networks, which visually match the anterior Saliency and left and right executive control network of the Stanford Template. Those matching RSNs were considered as corresponding templates and named according to the Stanford template anterior Saliency (aSN, matching the executive control of the Smith template), left and right executive control (LECN and RECN, respectively, matching the left and right frontoparietal network of the Smith template).

Similarity between Smith templates and 3T ICA components was assessed by spatial correlation of the zscore values of all voxels within the brain. Stanford templates were compared with the binarized ICA aggregate components (zscore > 2, corresponding to p<0.05, uncorrected) by spatial overlap with reference to the template and additionally the similarity was determined using the Jacquard index. RSNs were automatically identified by the maximum similarity converging in both directions (best match of all ICA components to a specific template and best match of all templates to specific ICA component) for either the Smith or the Stanford template. Subsequently the 7T RSNs were identified by spatial cross correlation of the 20 7T ICA components to the previously defined 3T RSNs. All automatically identified RSNs were confirmed by visual inspection (see Extended Data Fig. 4 and Supplementary results for details).

### Graph-theoretical resting-state analysis using multi seed correlation (MSRA)

According to the ICA analysis, the residuals of the preprocessed data after regression of white matter and ventricle time courses were used. Graph-theoretical resting-state analysis describes brain regions as nodes and focusses on the connections between each pair of brain region, called edges in network terminology. To assess the functional connectivity between the brain regions we used a multi-seed-region approach (MSRA) introduced by Kreitz et al. (2018)^27^. Briefly, a predefined seed region was placed automatically in the center of mass of each brain region as determined by atlas registration and region segmentation described above. Seed regions were spheres with approximately 7.5 mm diameter. Due to the different voxel resolution of 7T and 3T measurements odd kernel sizes to achieve closest diameters were 5 voxel for 7T (7.5 mm diameter) and 3 voxel for 3T (6 mm diameter). The average time course of each seed region was correlated with every voxel time course within the brain and the resulting correlation maps were thresholded using BY-FDR (q=0.05, n=298 time points) to determine significantly correlating voxels. In opposite to the predefined seed regions the location of those target voxels per brain region was determined purely data driven, thereby enhancing sensitivity of the connectivity between pairs of nodes^27^. The average Pearson’s correlation r of all target voxels per brain region was used to define the connectivity strength to the respective seed region. This procedure was repeated for every brain region resulting in an asymmetric correlation matrix per resting-state scan. For further analysis, Pearson’s r-values were transformed to Fisher’s z-values to provide normal distribution.

### Resting-state quality measures

Quality of resting-state connectivity was assessed via the reasonable assumption that homotopic brain regions in healthy subjects are stronger connected than heterotopic, independently of their anatomical distance^28^. For each bilateral brain region, the normalized rank of the connectivity strength to its counterpart in the opposite hemisphere within all its connections was determined. Subsequently, a linear fit over the ranks of all brain regions in dependence on the anatomical distance was calculated. Group comparisons between 7T and 3T resting-state matrices were conducted with the fitted rank for the mean anatomical distance and the slope of the linear fit. The first indicates the general dominance of bilateral interhemispheric connectivity and the latter the dependency on anatomical distance.

Another measure for resting-state data quality is the specificity for distinct RSNs. In theory, two core regions of the default mode network should correlate stronger than on of these regions to a core region of another, preferably task positive network^29^. Here, we calculated the correlation ratio of the anterior cingulate cortex to the middle cingulum (part of the central axis of the default mode network) and to the middle frontal gyrus (core region of the executive control network). Correlation of all hemispherical combinations were averaged (left to left, right to right, left to right and right to left). Higher values indicate higher specificity.

### Topological comparison of resting-state graphs

For topological comparison, the 7% strongest connections of the average matrices per group (7T and 3T) were extracted to create average networks of the same density. Network communities were detected using a heuristic method that is based on modularity optimization proposed by Blondel et al., (2008)^67^. The nodes within these communities are more strongly connected to each other than to nodes outside the community. Networks were visualized in AMIRA (Thermo Fisher Scientific Inc., Waltham, MA, USA, V5.4.2) using a force-based algorithm^68^.

Topological components that represent subnetworks of altered connectivity strength between the 7T rs1 and the 3T rs1 scan were determined using an adaptation of the network based statistics (NBS)^69^. NBS is a method to control the family-wise error rate after mass univariate t-tests performed at every single network edge. NBS exploits the interconnected extent of univariate significant different edges by permutation of subject specific networks between experimental groups. In a paired design, we introduced an additional paired control group in order to control for general effects of repeated measurements (pNBS)^27^. Here, we used an unpublished control data set of 11 healthy subjects (7 females, age 25 to 63) who were measured twice with an interval of two days on the same Siemens TRIO Magnetom Scanner (GE-EPI, TR: 3 sec, TE: 30 ms, flip angle: 90°, matrix size 128 × 128 pixel, pixel resolution: 1.5 mm × 1.5 mm, 36 slices, slice thickness: 3 mm, slice gap: 0.75 mm, 200 volumes). Preprocessing and MSRA analysis were performed as described above. This control group was used to define an α-value where almost no significantly different connections between the repeated measurements occurred. To minimize the effect of single outliers, we calculated the 99% quantile of the mass univariate paired t-statistics p-values. The resulting α-value was used to identify a set of supra-threshold connections (control component). The same threshold was applied to the paired t-statistics p-values of the experimental group (i.e. paired rs1 scans with 7T and 3T), and all remaining connected components equal or smaller to the control component were eliminated. Finally, the family-wise-error (p_FWE_) was controlled by 10,000 randomized permutations of pairs between experimental and control group (for details see^27^). The α-value ensures that the observed differences correspond dominantly to the experimental conditions, in this case the field strength, whereas the p_FWE_ value indicates the probability that the observed component is not random. NBS and pNBS are weak controls, which only allows rejecting the global null hypothesis. Thus, no single connections but rather the whole component mirrors the resting-state modulations under the experimental conditions.

### Variability and Reproducibility of resting-state correlation matrices

Variability of resting-state graphs between subjects was assessed via pairwise spatial correlation of the underlying MSRA matrices, resulting in a correlation matrix which represents the similarity of each subject with all other subjects. Reproducibility of two subsequent resting-state measurements was determined by spatial correlation of the MSRA matrices of rs1 and rs2 separately for each subject. Here, only subjects without finger-tapping stimulation between both resting-state measurements were taken into account. Correlation values were transformed into Fisher’s z-values.

### Analysis of Resting-state Modulations

Since we expect only tiny modulations in distinct brain regions, pNBS was not powerful enough to detect more modulations between rs1 and rs2 in the finger-tapping group compared to the rest-group. However, reinforced by the observed reduced variability between subjects in rs2 (see Results), the classical NBS (α=0.05, 10,000 permutations) had enough power to detect significant components in rs2 that distinguish between groups. This first step of the resting-state modulation analysis was completely data driven and was applied to rs1 and rs2, respectively.

The dominant regions within the significant component resulting from the rs2 group comparison, i.e. those with the most modulated connections, were used for further analysis. This second step should identify specific connections that are significantly modulated due to the finger-tapping task performed between both resting-state measurements. For this, we used the region specific seed correlation maps (SCM) calculated during the MSRA procedure for rs1 and rs2 in both groups. The MSRA algorithm ensured that the seed regions were subject- and measurement-specific placed according to their relative position within the brain region of interest and sized. Only voxels with significant correlation of their time courses to the seed region time course as identified using BY-FDR were taken into account. SCMs of the selected regions were registered to the MNI space by applying the transformation resulting from the diffeomorphic registration of their corresponding anatomical references (see “Individual brain atlas registration and brain region segmentation”).

Subsequently, a voxel-wise paired t-test between rs1 and rs2 was performed separately for each group. The resulting statistical t-maps was cluster enhanced using threshold-free cluster enhancement (TFCE with parameter settings signal height H=2 and cluster extent E=0.05 ^70^) and corrected for multiple comparison using permutation testing. Relying on the assumption that significance of a contiguous cluster is more likely to be true positive than that of a single voxel, TFCE aims to enhance areas of t-values that exhibit some spatial contiguity without the need for a hard cluster-forming thresholding. The resulting modified values are normalized back to the range of the original t-values. Randomized permutation testing was performed with 1,000 repetitions. To avoid outliers and therefore improve statistical power the 99.9^th^ percentile of all voxel t-values within the brain instead of the maximum t-value was used to create the “Null”-distribution. The 95% quantile of this distribution represented the rejection condition and the normalized TFCE image was thresholded with this t-value to obtain significant clusters. The combination of both methods provided enhanced sensitivity (less false negative, reduced Type II error) of statistical tests on behalf of acceptable costs in specificity (more false positive, enhanced Type I error). Binarized significant clusters of the ft- and the rest-group were combined by a logical OR. This resulted in a binary mask that marked all voxels that are significant in either of both groups. This mask was applied to each subject’s rs2 – rs1 difference map and, after brain region segmentation using the maximum probability map of the above-described 4D probability atlas in MNI space, the average difference per subject and brain region was extracted.

To obtain specific connectivity modulations due to finger-tapping, the rest-group serves as a control and region specific average rs2-rs1 differences were tested for significance between ft- and rest-group using a homoscedastic t-test and Benjamini-Hochberg FDR^71^ (BH-FDR, q=0.05) to correct for multiple comparison. The resulting connections were visualized as network graphs with brain regions as nodes and the modified connections of the seed regions as edges using AMIRA (Thermo Fisher Scientific Inc., Waltham, MA, USA, V5.4.2).

### Statistics

In addition to network based statistics (NBS and pNBS) and voxel wise paired t-test including TFCE, as described above, the following statistical approaches were used:

Principle Component Analysis was performed to separate 3T and 7T measurements by quality metrics (n=18) and by BOLD response parameters including tCNR (n=9). To search for group specific BOLD response parameter effects, we performed a mixed repeated measure ANOVA with replication using field strength as within factor and brain regions as between factor (n=9). Group effects of the activation probability were assessed using two factor ANOVA without replication with factors field strength and brain regions. Variability main effects between measurements were examined by one factor repeated measure ANOVA using Fisher’s z correlation values of all subject pairs as a measure of similarity (n=36). Significant ANOVA effects and interactions were then tested using Tukey HSD. Direct two group comparisons were made using Student’s t-test, either paired (3T vs 7T, rs1 vs rs2) or homoscedastic (ft vs rest). Multiple tests were appropriately corrected using permutation tests (NBS, pNBS, voxel wise t-statistic, description see above), Benjamini-Yekutieli FDR (q=0.05, detection of significant BOLD response and resting-state correlation in volumes with voxel wise statistic), Benjamini-Hochberg FDR (q=0.05, detection of significant group differences in region specific statistics) and Bonferroni (correction of ANOVA follow up Tukey HSD statistics). Significance level was p<0.05.

### Data availability

The data that support the findings of this study are available from the corresponding author upon reasonable request. Template sets of Resting-State Networks used in this study are available from https://www.fmrib.ox.ac.uk/datasets/brainmap+rsns/ and http://findlab.stanford.edu/functional_ROIs.html

### Code availability

fMRI data were preprocessed using Brainvoyager QX 2.8.2.2523 (Brain Innovation, Maastricht, Netherlands), Data analysis was conducted in Magnan 2.5 (BioCom GbR, Uttenreuth, Germany), image registration to MNI space was performed using Advanced Normalization Tools (ANTS55, http://stnava.github.io/ANTs/), and ICA was done by GIFT v1.3g (https://trendscenter.org/software/). For visulization we used AMIRA (Thermo Fisher Scientific Inc., Waltham, MA, USA, V5.4.2), and Microsoft Excel 2016 including the freely available software package Real Statistics (https://www.real-statistics.com/) for further statistical analysis.

## Supporting information

Supplemental Results and Discussion

Tables 1 and 3: PCA loadings

## Extended Data

**Extended Data Fig. 1:**
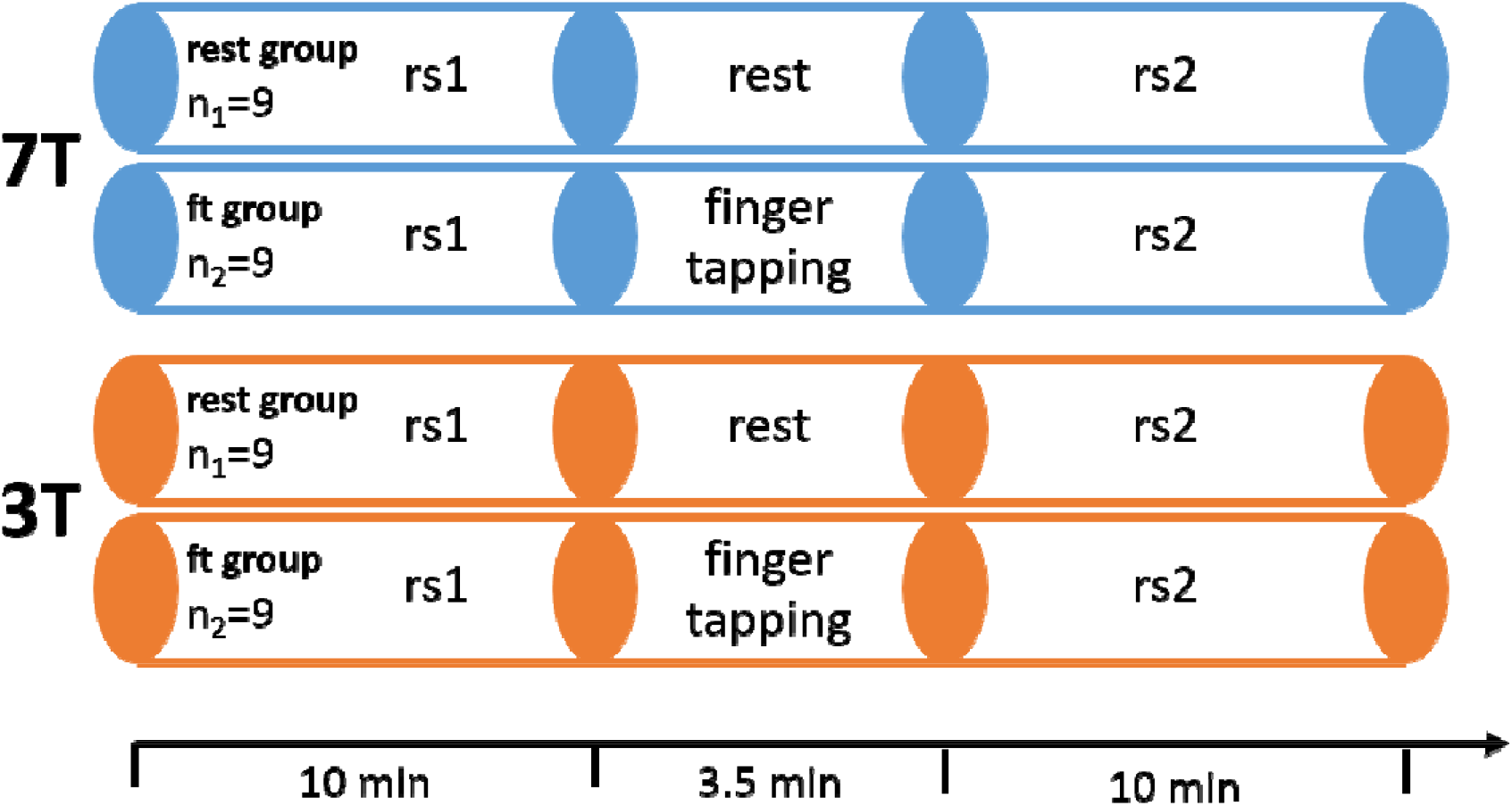
Experimental design. 18 healthy participants were measured twice, once in a 7T and once in a 3T Siemens Magnetom Scanner in a randomized order. Each session consisted of two resting state scans (rs1 and rs2) with either rest or a simple finger-tapping motor task in between. In the active motor task, participants tapped each finger sequentially for 14 seconds, followed by 14 seconds of rest, for a total of 7 times after an initial 14 seconds rest (starting and stopping upon voice commands).

**Extended Data Fig. 2:**
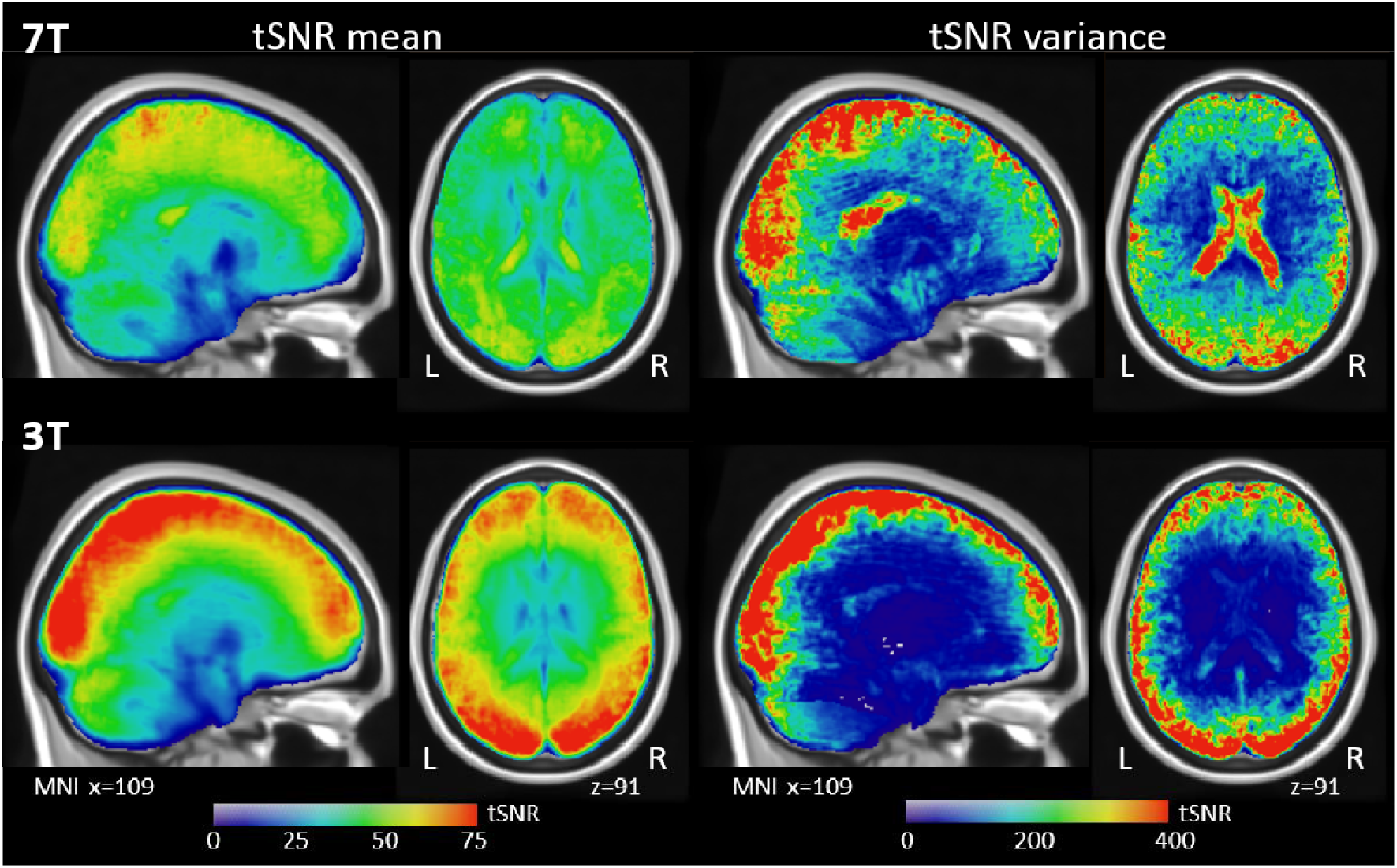
Spatial distribution of tSNR throughout the brain for the first resting-state. Top: 7T rs1 scan, bottom: 3T rs1 scan. Average (left) and variance (right) heat maps are superimposed on MNI standard space template. L: left hemisphere, R: right hemisphere.

**Extended Data Fig. 3:**
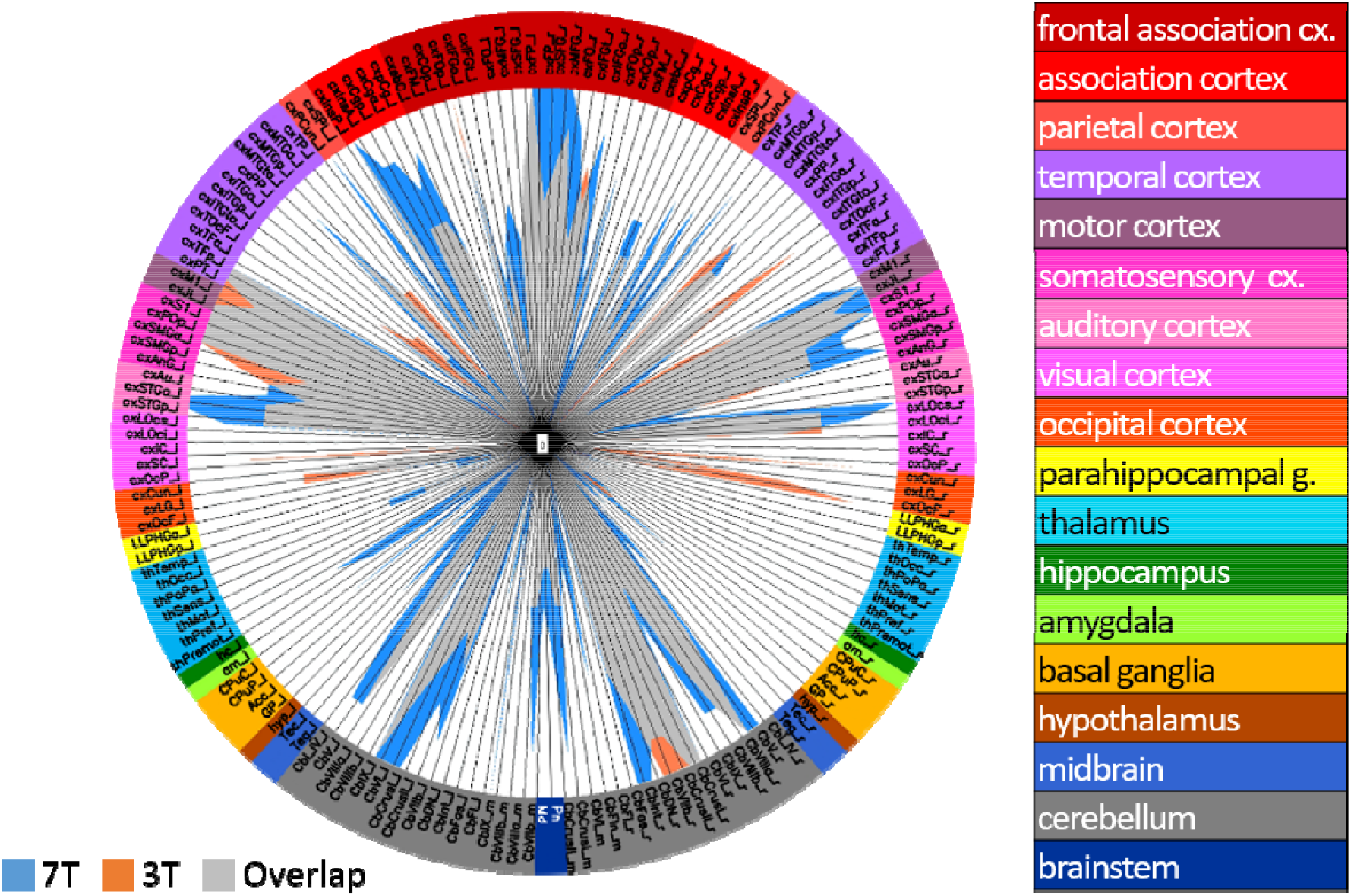
Regional activation propability due to a finger-tapping motor task. Each participant was scanned with 7T and 3T, respectively. The activation propability is the proportion of participants that show motor task induced activation in a respective brain region.

**Extended Data Fig. 4:**
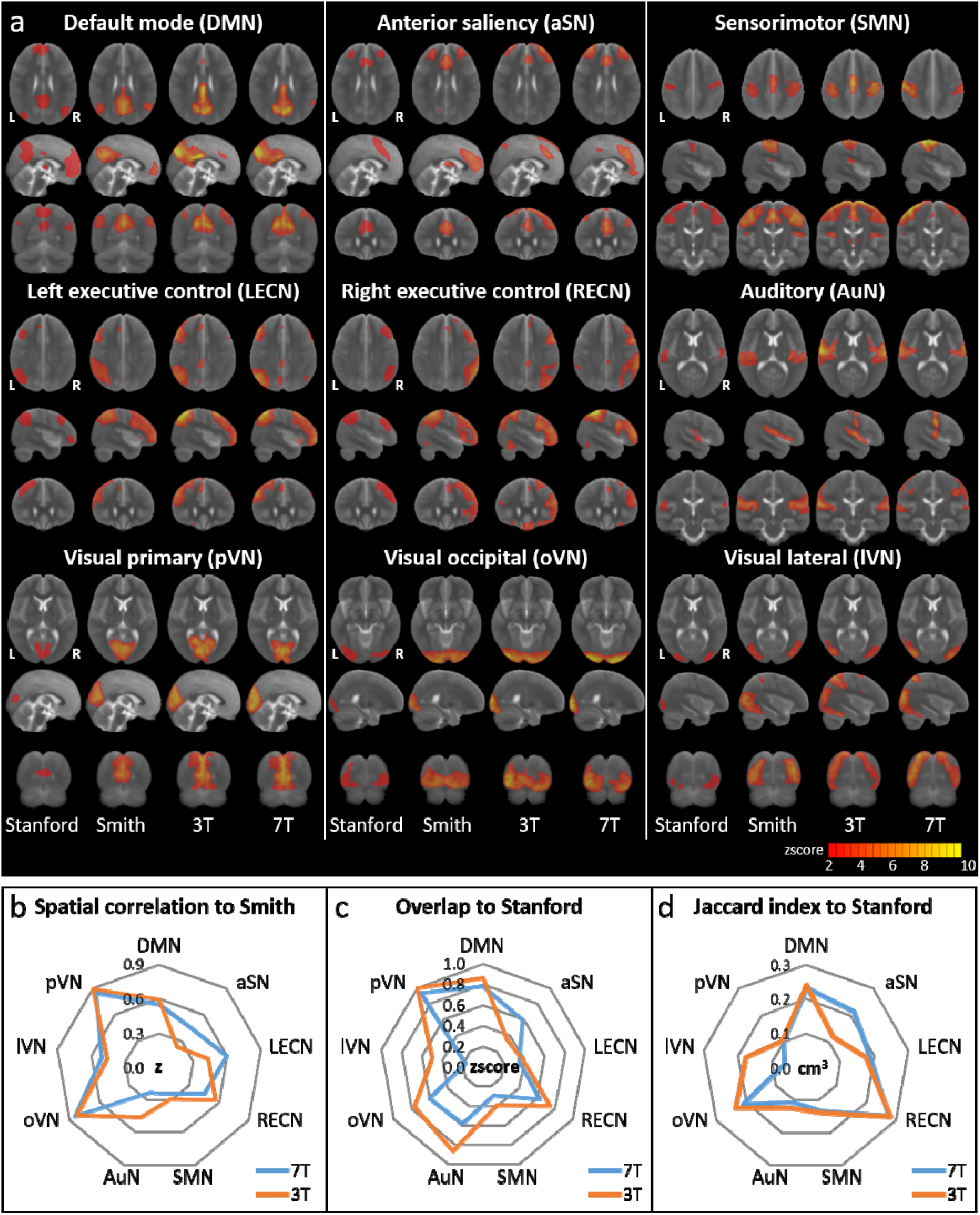
Identification of resting-state networks derived from 20 component group ICA. a) Public available resting-state network (RSN) templates (first column of each set: Stanford, second column: Smith, for details see Methods) and goup ICA aggregate components (zscore maps) of 3T (third column) and 7T (fourth column) resting-state rs1 scans. 3T and 7T zscore maps were thresholded at zscore=2 (corresponding to p<0.05, uncorrected). The Smith template was thresholded accordingly to match the 3T zscore map best. The Stanford template is provided binarized. The 3 most informative orthogonal slices for each set and RSN was visualized superimposed on the MNI standard space template image. b) Identification of RSNs via spatial cross correlation of 3T and 7T ICA zscore maps to the Smith template. c) Identification of RSNs via spatial overlap of thresholded (zscore > 2) and binarized 3T and 7T ICA zscore maps with Stanford template. d) Similarity of thresholded (zscore > 2) and binarized 3T and 7T ICA zscore maps with Stanford template assessed via the jaccard index.

**Extended Data Fig. 5:**
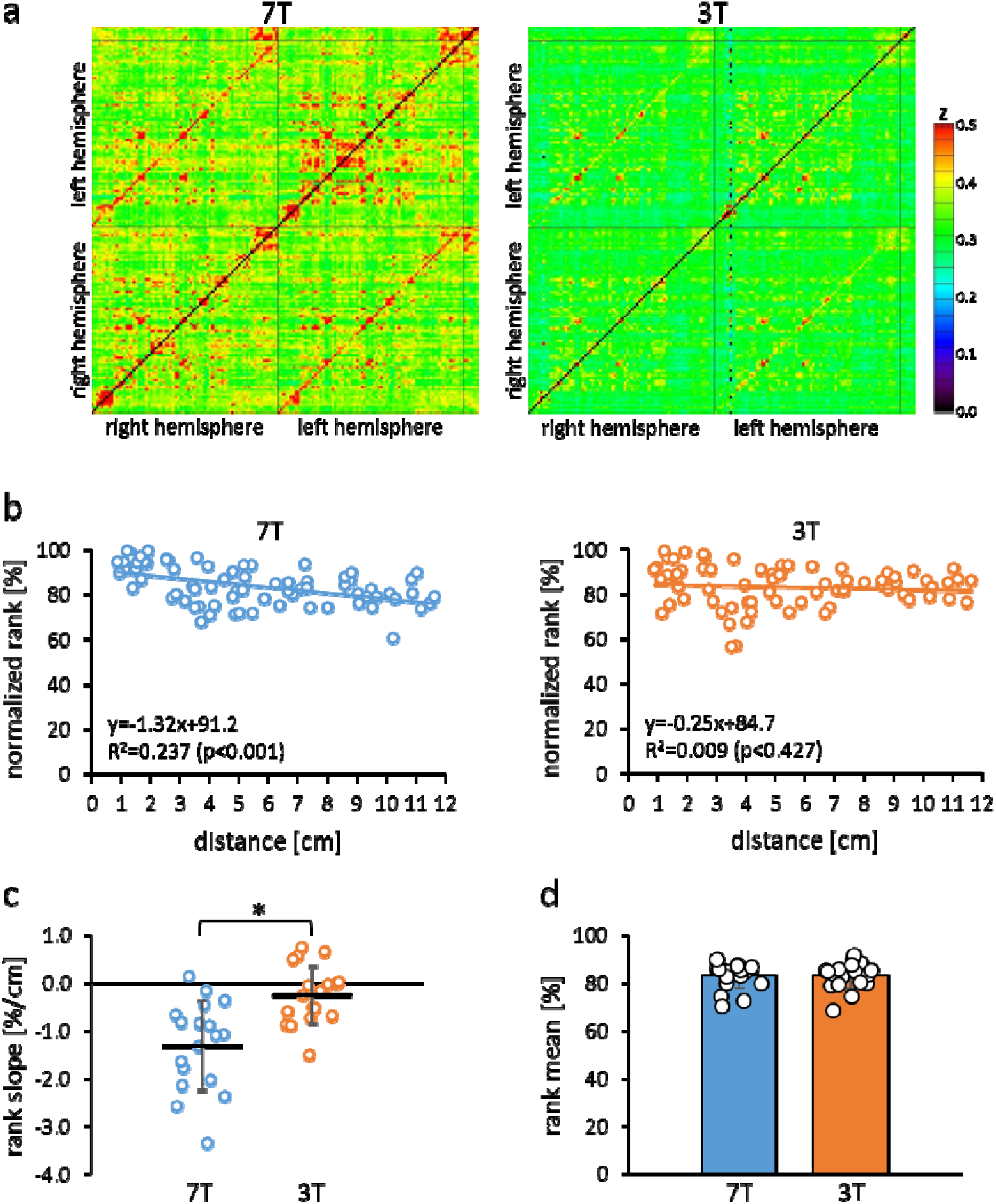
Quality assessment of resting state matrices derived from multi-seed-region analysis (MSRA). a) Average MSRA correlation matrices resulting from rs1 scans using 7T (left) and 3T (right) fMRI. b) Ranked average correlation of bilateral brain regions across hemispheres dependent on their anatomical distance per subject rs1 scan using 7T (left) and 3T (right) fMRI, including linear trendlines. c) Slope of the linear trendline described in b) per subject (n=18, *p < 0.05, paired t-test). d) Average correlation rank of all bilateral regions per subject. No significant differences between 7T and 3T were observed (n=18, paired t-test)

**Extended Data Fig. 6:**
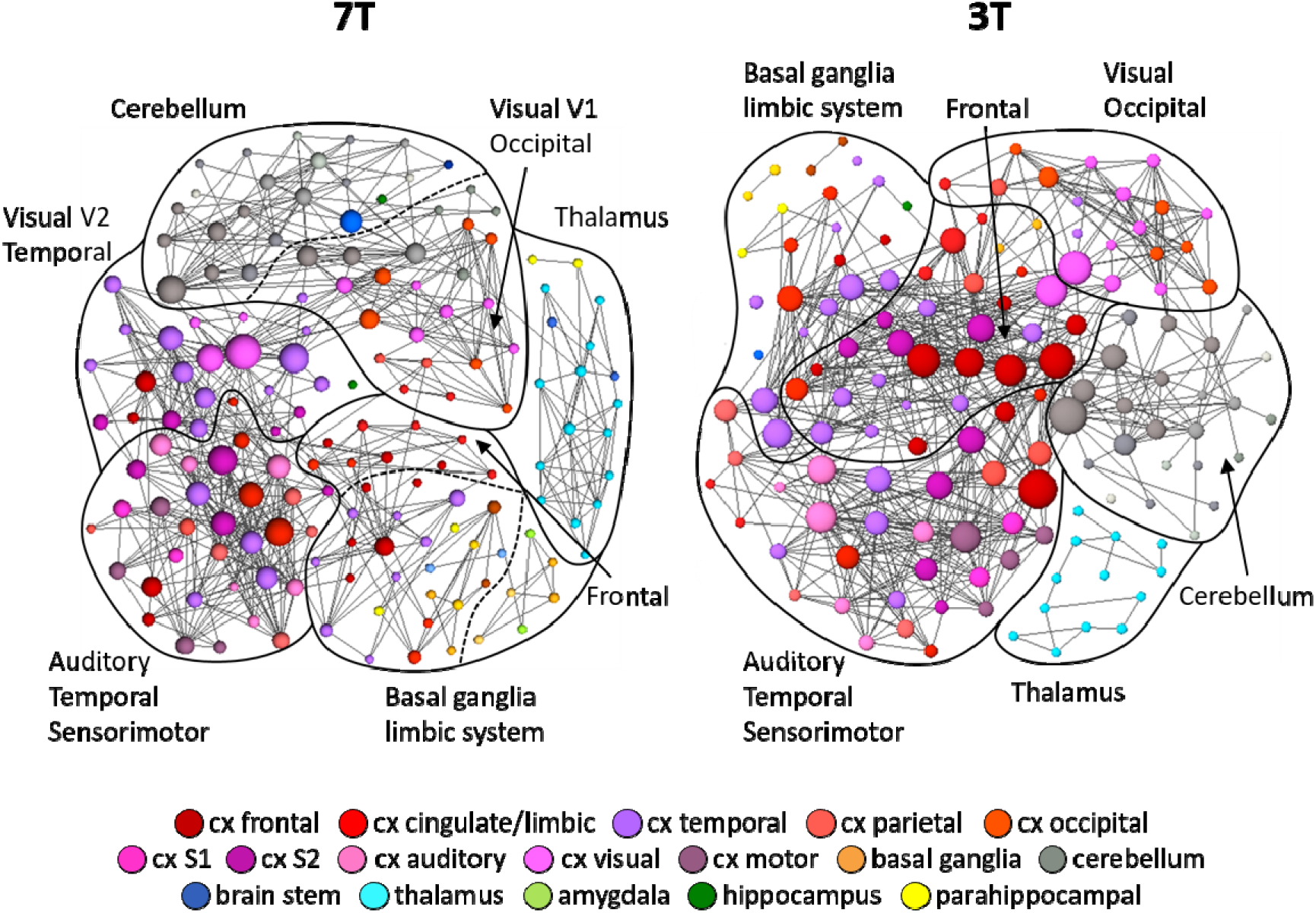
Communities of resting-state MSRA graphs. Average 7T (left) and 3T (right) first resting-state rs1 scans, normalized to a density of 7% of all possible connections are shown. Communities were detected using the Blondel algorithm^66^ and marked by surrounding solid (community level 1) and dashed (community level 0) lines. Node size represent their degree (i.e. the number of connections per node). Graphs were visualized using a forced based algorithm^67^.

**Extended Data Fig. 7:**
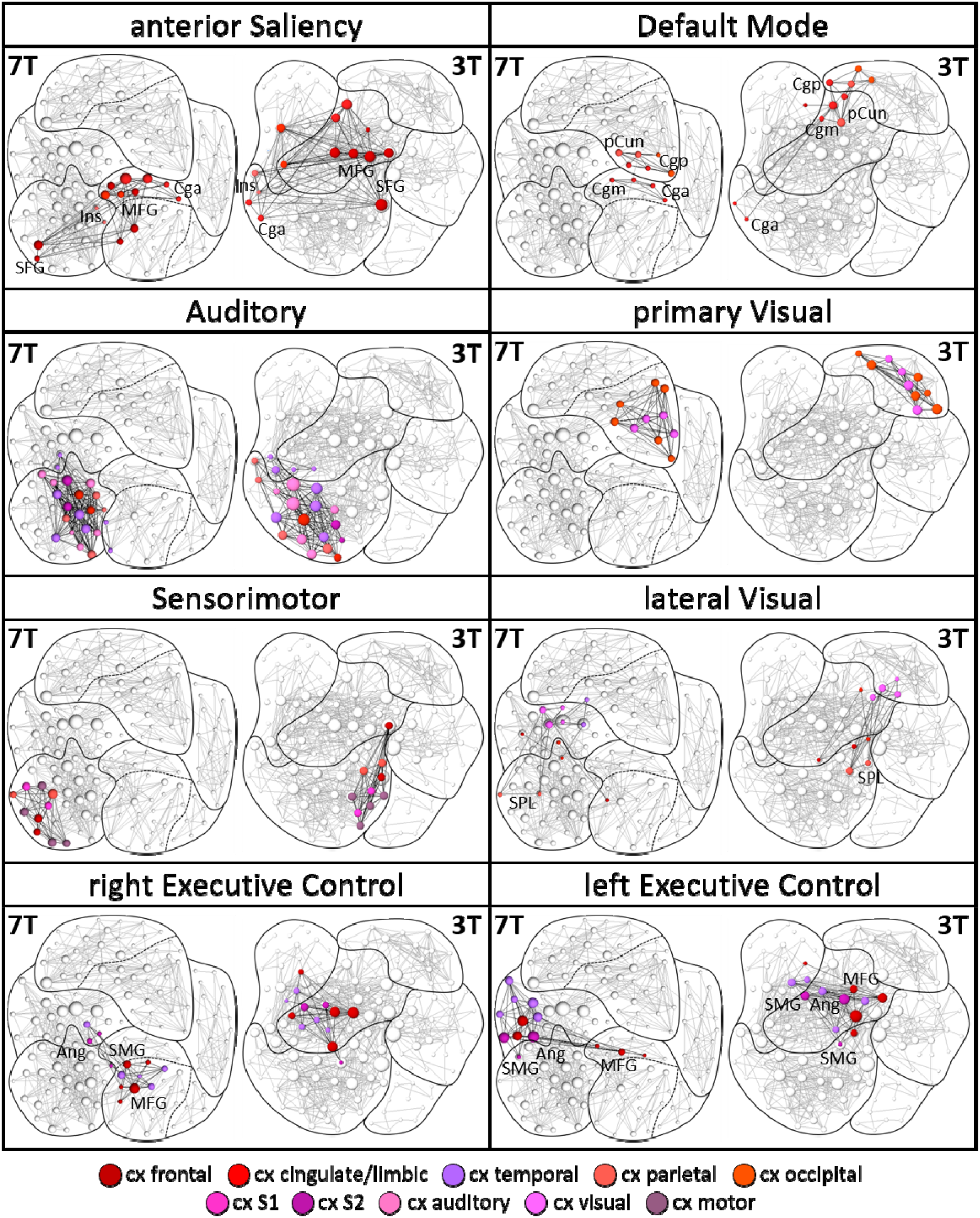
Brain regions contributing to RSNs derived from group ICA aggregate components presented as nodes superimposed on MSRA community graphs. Node size code for their degree in a complete graph normalized to 7% density. The occipital visual RSN largely overlaps with the primary visual RSN. The dominant region characterizing solely the occipital visual RSN is the occipital pole, located in the same community as and closely related to the primary visual RSN. Therefore, as it hardly represents a graph theoretical subnetwork, the occcipital RSN is not shown. Ang: angular gyrus, Cga: anterior cinculate cortex, Cgm: middle cingulate cortex. Cgp: posterior cingulate cortex, Ins: insula, MFG: middle frontal gyrus, pCun: precuneus, SMG: supramarginal gyrus, SFG: superior frontal gyrus.

**Extended Data Fig. 8:**
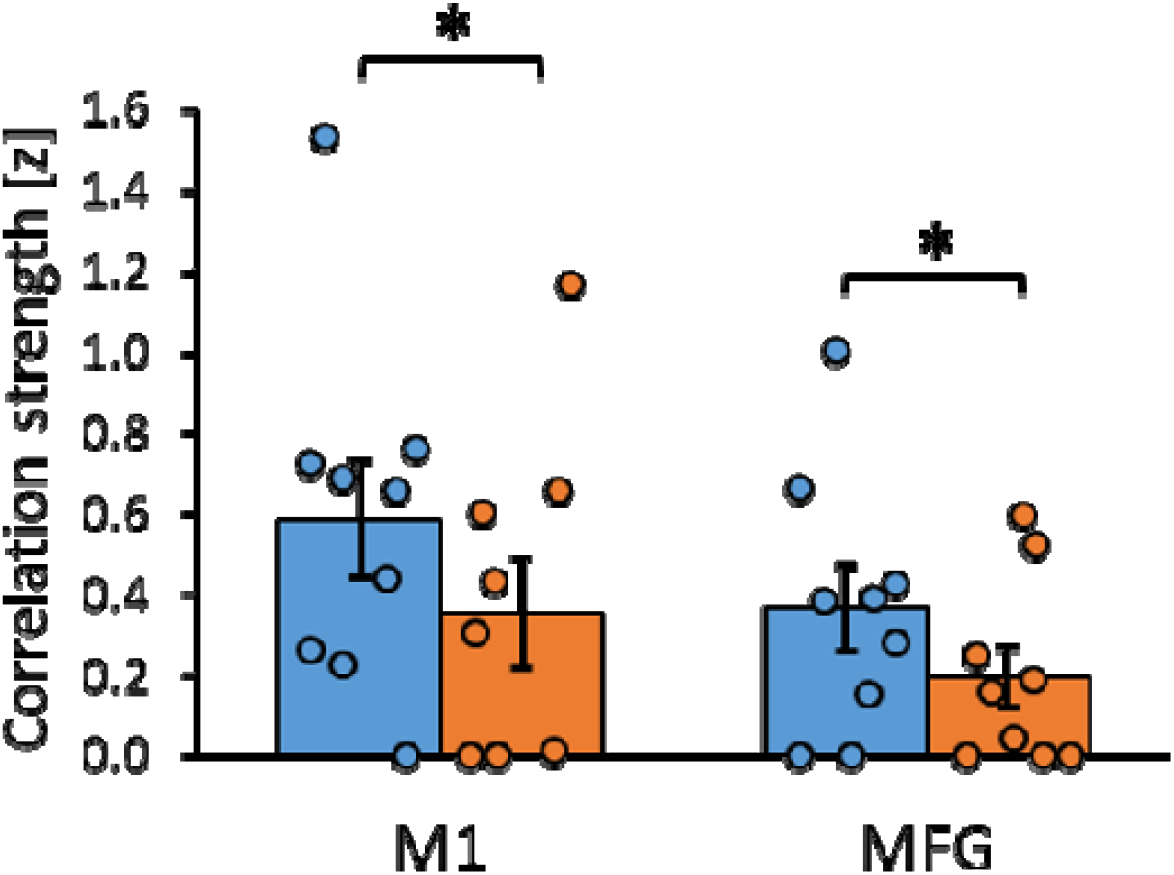
Interhemispheric functional connectivity during task performance. Functional connectivity was assessed via correlation of average regional time courses after regressing out the global mean time course of all activated regions to eliminate the stimulation driven response. * paired t-test, p<0.05, n=9

