## Supplemental Results and Discussion for "Capturing early human memory consolidation utilizing the higher functional specificity of 7T compared to 3T-fMRI"

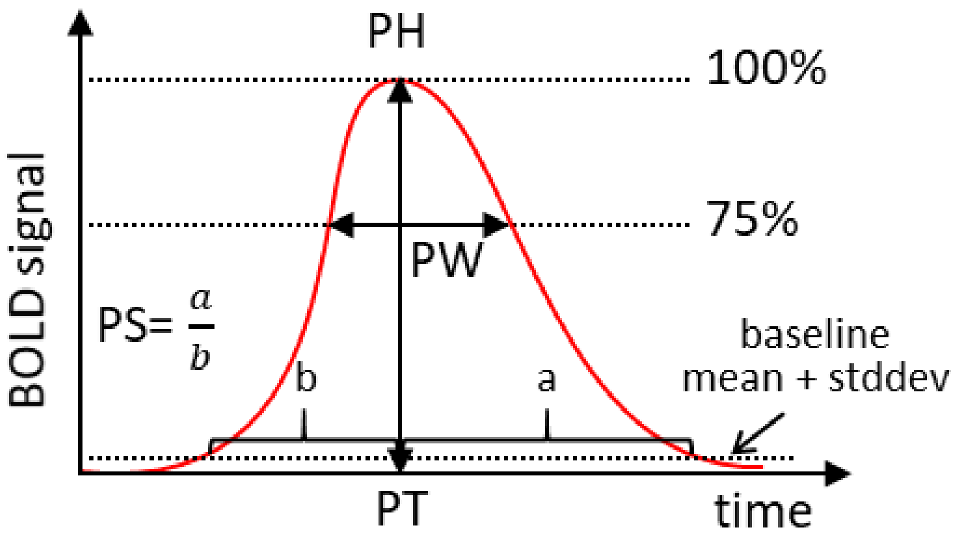


**Supplementary Fig. 1: BOLD response parameters describing the BOLD signal response amplitude.**

PH: amplitude (peak) height, PS: amplitude symmetry, PT: time to peak, PW: peakwidth. For more detailed description see online methods.

**Supplementary Table 1: PCA of quality metrics. Loadings of the first 5 PCs.**

| | Eigenvalue [%] | 38.8 | 60.4 | 76.8 | 87.9 | 95.9 |
|---|---|---|---|---|---|---|
| **Principle Component** | | **PC1** | **PC2** | **PC3** | **PC4** | **PC5** |
| temporal | tSNR | -0.288 | 0.258 | -0.572 | -0.087 | 0.479 |
| | zDVARS | -0.212 | 0.449 | -0.148 | -0.441 | -0.691 |
| | MDI | -0.376 | -0.271 | 0.451 | 0.268 | -0.235 |
| | Gcorr | 0.067 | -0.100 | -0.566 | 0.704 | -0.402 |
| spatial | SNR | -0.272 | 0.557 | 0.279 | 0.330 | 0.082 |
| | CNR | 0.509 | 0.195 | 0.030 | -0.041 | -0.214 |
| | FBER | 0.304 | 0.543 | 0.219 | 0.332 | 0.150 |
| | EFC | 0.548 | -0.070 | 0.011 | -0.100 | 0.029 |

Loadings above 0.5 for each PC are highlighted in grey. Only PC1 contribute to the separation between 7T and 3T measurements (indicated by the red frame).

**Supplementary Table 2: BOLD response parameters to finger tapping motor stimulation obtained with 7T and 3T fMRI**

| Anatomical group | Abbr. | Area | 7 T | | | | | | | | | | | | 3T | | | | | | | | | | | |
|---|---|---|---|---|---|---|---|---|---|---|---|---|---|---|---|---|---|---|---|---|---|---|---|---|---|---|
| | | | Activation propability [%] | | Activated volume [cm$^3$] | | Amplitude peak height [% signal] | | Amplitude width [sec] | | Amplitude symmetry [au] | | Amplitude peak time [sec] | | activation propability [%] | | activated volume [cm$^3$] | | amplitude peak height [% signal] | | amplitude width [sec] | | amplitude symmetry [au] | | amplitude peak time [sec] | |
| | | | l | r | l | r | l | r | l | r | l | r | l | r | l | r | l | r | l | r | l | r | l | r | l | r |
| primary somato-sensory cortex | cxS1 | Postcentral Gyrus | 88.9 | 100.0 | 7.02 (0.83) | 3.44 (0.92) | 2.47 (0.23) | 1.22 (0.09) | 21.40 (1.54) | 19.52 (1.47) | 1.51 (0.41) | 1.49 (0.41) | 8.67 (1.05) | 9.33 (1.25) | 66.7 | 100.0 | 7.23 (0.67) | 2.80 (0.89) | 1.30 (0.13) | 0.83 (0.11) | 19.52 (1.47) | 19.34 (1.47) | 1.14 (0.36) | 1.60 (0.42) | 11.11 (0.59) | 9.56 (0.44) |
| secondary somatosensory cortex | cxPOp | Parietal Operculum Cortex | 77.8 | 88.9 | 0.40 (0.08) | 0.22 (0.05) | 1.21 (0.15) | 0.86 (0.21) | 19.69 (1.48) | 18.47 (1.43) | 1.50 (0.41) | 1.46 (0.40) | 10.00 (1.53) | 10.33 (1.41) | 77.8 | 100.0 | 0.35 (0.13) | 0.17 (0.08) | 0.69 (0.12) | 0.56 (0.13) | 20.04 (1.49) | 19.81 (1.48) | 1.58 (0.42) | 1.35 (0.39) | 10.00 (0.75) | 10.56 (0.93) |
| | cxSMGa | Supramarginal Gyrus, anterior | 100.0 | 100.0 | 2.33 (0.34) | 0.95 (0.24) | 1.63 (0.16) | 0.94 (0.16) | 20.65 (1.51) | 20.19 (1.50) | 1.53 (0.41) | 1.98 (0.47) | 9.33 (1.15) | 9.22 (1.37) | 100.0 | 66.7 | 1.94 (0.41) | 0.67 (0.36) | 1.00 (0.13) | 0.48 (0.13) | 19.02 (1.45) | 19.08 (1.46) | 1.37 (0.39) | 1.63 (0.43) | 10.67 (0.67) | 9.33 (0.44) |
| | cxSMGp | Supramarginal Gyrus, posterior | 100.0 | 100.0 | 1.31 (0.25) | 0.52 (0.12) | 1.50 (0.22) | 0.99 (0.05) | 20.77 (1.52) | 19.54 (1.47) | 1.64 (0.43) | 1.43 (0.4) | 9.33 (1.45) | 10.89 (1.38) | 100.0 | 100.0 | 1.12 (0.27) | 0.35 (0.12) | 0.88 (0.14) | 0.63 (0.14) | 20.12 (1.50) | 19.71 (1.48) | 1.79 (0.45) | 1.69 (0.43) | 9.67 (0.58) | 9.33 (0.55) |
| auditory cortex | cxSTGp | Superior Temporal Gyrus, posterior | 88.9 | 100.0 | 0.42 (0.13) | 0.46 (0.16) | 1.59 (0.15) | 1.46 (0.18) | 19.30 (1.46) | 18.59 (1.44) | 1.11 (0.35) | 1.73 (0.44) | 12.44 (1.44) | 9.78 (1.58) | 100.0 | 100.0 | 0.26 (0.11) | 0.41 (0.16) | 0.67 (0.19) | 0.68 (0.17) | 19.12 (1.46) | 19.22 (1.46) | 1.67 (0.43) | 1.34 (0.39) | 9.67 (0.55) | 9.89 (0.86) |
| motor cortex | cxM1 | Precentral Gyrus | 100.0 | 100.0 | 7.47 (0.85) | 3.18 (0.99) | 2.48 (0.36) | 1.23 (0.13) | 21.70 (1.55) | 19.90 (1.49) | 1.50 (0.41) | 1.97 (0.47) | 8.89 (1.16) | 8.22 (1.35) | 100.0 | 100.0 | 7.62 (0.98) | 2.93 (0.96) | 1.37 (0.14) | 0.83 (0.09) | 19.21 (1.46) | 19.07 (1.46) | 1.21 (0.37) | 1.33 (0.38) | 10.89 (0.48) | 9.56 (0.44) |
| | cxJL | Juxtapositional Lobule Cortex | 100.0 | 100.0 | 1.51 (0.27) | 0.95 (0.22) | 1.58 (0.23) | 1.43 (0.21) | 20.20 (1.50) | 19.78 (1.48) | 2.00 (0.47) | 1.96 (0.47) | 8.33 (1.35) | 8.78 (1.41) | 100.0 | 77.8 | 1.42 (0.28) | 0.86 (0.27) | 1.12 (0.13) | 1.02 (0.14) | 19.68 (1.48) | 19.57 (1.47) | 1.68 (0.43) | 1.58 (0.42) | 8.89 (0.48) | 9.56 (0.87) |
| temporal cortex | cxTP | Temporal Pole | 66.7 | 88.9 | 0.62 (0.17) | 0.77 (0.28) | 1.39 (0.14) | 1.30 (0.22) | 18.54 (1.44) | 19.43 (1.47) | 1.74 (0.44) | 1.72 (0.44) | 10.67 (1.56) | 11.00 (1.62) | 33.3 | 55.6 | 0.25 (0.12) | 0.50 (0.19) | 0.43 (0.18) | 1.12 (0.18) | 19.90 (1.49) | 19.29 (1.46) | 1.46 (0.40) | 1.61 (0.42) | 10.11 (0.56) | 10.11 (0.79) |
| | cxPT | Planum Temporale | 100.0 | 100.0 | 0.67 (0.14) | 0.20 (0.05) | 1.39 (0.12) | 1.12 (0.20) | 19.80 (1.48) | 19.14 (1.46) | 1.53 (0.41) | 1.04 (0.34) | 10.44 (1.59) | 11.78 (1.47) | 100.0 | 77.8 | 0.42 (0.11) | 0.20 (0.07) | 0.78 (0.05) | 0.58 (0.17) | 20.13 (1.5) | 18.94 (1.45) | 1.47 (0.40) | 1.41 (0.40) | 10.44 (0.80) | 9.78 (0.86) |
| parietal cortex | cxSPL | Superior Parietal Lobule | 100.0 | 77.8 | 2.26 (0.47) | 1.99 (0.56) | 1.59 (0.20) | 0.90 (0.19) | 19.26 (1.46) | 18.85 (1.45) | 1.94 (0.46) | 2.39 (0.52) | 9.11 (1.38) | 9.00 (1.37) | 66.7 | 55.6 | 1.95 (0.60) | 1.67 (0.76) | 0.83 (0.09) | 0.53 (0.16) | 19.66 (1.48) | 19.57 (1.47) | 1.38 (0.39) | 1.45 (0.40) | 10.22 (0.85) | 9.89 (0.65) |
| frontal cortex | cxCOp | Central Opercular Cortex | 88.9 | 100.0 | 0.68 (0.14) | 0.38 (0.09) | 0.91 (0.14) | 0.83 (0.16) | 21.43 (1.54) | 21.42 (1.54) | 1.48 (0.41) | 1.30 (0.38) | 9.44 (1.32) | 9.67 (1.18) | 66.7 | 88.9 | 0.55 (0.18) | 0.41 (0.15) | 0.71 (0.05) | 0.54 (0.11) | 19.96 (1.49) | 19.88 (1.49) | 1.53 (0.41) | 1.30 (0.38) | 10.00 (0.58) | 10.22 (0.86) |
| | cxMFG | Middle Frontal Gyrus | 100.0 | 88.9 | 0.74 (0.40) | 0.73 (0.26) | 1.41 (0.41) | 1.36 (0.24) | 19.03 (1.45) | 18.31 (1.43) | 1.36 (0.39) | 2.13 (0.49) | 9.89 (1.16) | 9.56 (1.48) | 77.8 | 100.0 | 0.74 (0.31) | 0.96 (0.35) | 1.01 (0.25) | 0.85 (0.21) | 18.68 (1.44) | 18.93 (1.45) | 1.35 (0.39) | 1.42 (0.40) | 11.11 (0.51) | 9.67 (0.53) |
| | cxSFG | Superior Frontal Gyrus | 88.9 | 100.0 | 1.75 (0.46) | 1.31 (0.40) | 2.01 (0.28) | 1.23 (0.11) | 18.97 (1.45) | 19.15 (1.46) | 1.63 (0.43) | 1.78 (0.44) | 10.44 (1.41) | 11.11 (1.74) | 100.0 | 100.0 | 1.59 (0.47) | 1.36 (0.63) | 1.37 (0.28) | 0.86 (0.25) | 20.19 (1.50) | 19.20 (1.46) | 1.50 (0.41) | 1.58 (0.42) | 11.00 (0.97) | 9.56 (0.67) |
| | cxFP | Frontal Pole | 100.0 | 100.0 | 0.83 (0.26) | 1.10 (0.38) | 0.73 (0.14) | 0.79 (0.08) | 17.50 (1.39) | 19.95 (1.49) | 1.52 (0.41) | 1.09 (0.35) | 10.11 (1.14) | 11.78 (1.13) | 100.0 | 100.0 | 0.37 (0.17) | 0.72 (0.39) | 0.32 (0.11) | 0.29 (0.12) | 18.91 (1.45) | 18.80 (1.45) | 1.32 (0.38) | 1.16 (0.36) | 10.11 (0.73) | 10.22 (0.57) |
| cerebellum anterior lobe | CbI_IV | Area I_IV | 100.0 | 100.0 | 0.20 (0.12) | 0.50 (0.13) | 0.91 (0.27) | 1.21 (0.18) | 18.64 (1.44) | 18.57 (1.44) | 1.37 (0.39) | 1.90 (0.46) | 10.33 (1.33) | 9.78 (1.22) | 66.7 | 77.8 | 0.10 (0.06) | 0.29 (0.14) | 0.25 (0.13) | 0.33 (0.17) | 18.88 (1.45) | 18.99 (1.45) | 1.30 (0.38) | 1.07 (0.34) | 10.33 (0.53) | 10.78 (0.55) |
| | CbV | Area V | 100.0 | 88.9 | 0.32 (0.13) | 1.06 (0.12) | 0.95 (0.18) | 1.42 (0.20) | 19.50 (1.47) | 21.25 (1.54) | 1.95 (0.46) | 1.31 (0.38) | 9.56 (1.44) | 10.22 (0.97) | 77.8 | 77.8 | 0.35 (0.17) | 1.14 (0.29) | 0.45 (0.15) | 0.86 (0.13) | 19.32 (1.47) | 20.18 (1.50) | 1.37 (0.39) | 0.86 (0.31) | 10.56 (0.41) | 13.00 (0.33) |
| cerebellum superior posterior lobe | CbVI | Area VI | 88.9 | 100.0 | 1.52 (0.30) | 2.71 (0.36) | 1.03 (0.09) | 1.43 (0.09) | 20.32 (1.50) | 20.73 (1.52) | 0.83 (0.3) | 0.90 (0.32) | 12.00 (0.75) | 11.78 (0.85) | 77.8 | 100.0 | 1.51 (0.57) | 3.18 (0.40) | 0.81 (0.10) | 0.95 (0.06) | 18.13 (1.42) | 19.18 (1.46) | 1.57 (0.42) | 1.41 (0.40) | 10.44 (0.73) | 10.89 (0.68) |
| | CbCrusI | Area CrusI | 88.9 | 100.0 | 1.34 (0.20) | 1.70 (0.31) | 0.97 (0.06) | 1.16 (0.10) | 20.07 (1.49) | 18.81 (1.45) | 1.58 (0.42) | 1.94 (0.46) | 10.67 (1.29) | 9.56 (1.14) | 100.0 | 100.0 | 1.04 (0.47) | 1.07 (0.43) | 0.65 (0.16) | 0.84 (0.17) | 18.48 (1.43) | 17.41 (1.39) | 1.44 (0.40) | 1.32 (0.38) | 10.22 (0.55) | 10.67 (0.47) |
| | CbCrusII | Area CrusII | 88.9 | 88.9 | 0.21 (0.06) | 0.26 (0.07) | 0.82 (0.06) | 1.05 (0.19) | 18.82 (1.45) | 19.18 (1.46) | 0.98 (0.33) | 1.83 (0.45) | 12.67 (1.00) | 9.67 (1.37) | 66.7 | 88.9 | 0.40 (0.18) | 0.57 (0.26) | 0.73 (0.22) | 0.86 (0.15) | 18.61 (1.44) | 18.48 (1.43) | 1.57 (0.42) | 1.42 (0.40) | 10.11 (0.51) | 10.78 (0.70) |
| | CbVIIb | Area VIIb | 77.8 | 77.8 | 0.27 (0.11) | 0.60 (0.15) | 0.68 (0.14) | 0.87 (0.13) | 18.06 (1.42) | 21.45 (1.54) | 0.62 (0.26) | 1.35 (0.39) | 12.89 (0.87) | 10.33 (1.15) | 66.7 | 77.8 | 0.40 (0.19) | 0.69 (0.23) | 0.45 (0.16) | 0.75 (0.09) | 18.71 (1.44) | 18.58 (1.44) | 1.37 (0.39) | 1.50 (0.41) | 10.33 (0.55) | 10.67 (0.58) |
| cerebllum deep nuclei | CbDN | Dentate nuclei | 100.0 | 100.0 | 0.51 (0.18) | 1.12 (0.16) | 0.68 (0.10) | 1.28 (0.08) | 18.98 (1.45) | 20.34 (1.50) | 1.06 (0.34) | 1.25 (0.37) | 11.33 (1.11) | 11.11 (1.06) | 77.8 | 66.7 | 0.06 (0.03) | 0.15 (0.05) | 0.39 (0.11) | 0.69 (0.17) | 18.58 (1.44) | 18.54 (1.44) | 1.26 (0.37) | 1.29 (0.38) | 10.44 (0.58) | 10.89 (0.70) |
| cerebellum vermis | CbVI | Area VI m | 100.0 | | 0.51 (0.09) | | 1.36 (0.20) | | 20.67 (1.52) | | 2.22 (0.50) | | 10.00 (1.49) | | 88.9 | | 0.35 (0.11) | | 0.61 (0.13) | | 18.99 (1.45) | | 1.37 (0.39) | | 10.89 (0.90) | |
| **All** | | | **94.06 (1.28)** | | **57.02 (0.62)** | | **1.24 (0.22)** | | **19.67 (1.08)** | | **1.55 (0.47)** | | **10.22 (1.29)** | | **84.24 (2.54)** | | **51.12 (0.65)** | | **0.74 (0.17)** | | **19.2 (0.49)** | | **1.42 (0.20)** | | **10.28 (0.66)** | |

Values are means with standard errors in brackets. n=18, paired design. l: left hemisphere, r: right hemisphere

**Supplementary Table 3: PCA of BOLD response parameters. Loadings of the first 4 PCs.**

| Eigenvalue [%] | 33.0 | 65.0 | 86.0 | 96.7 |
|---|---|---|---|---|
| **Principle Component** | **PC1** | **PC2** | **PC3** | **PC4** |
| activation volume | 0.295 | -0.115 | -0.635 | -0.674 |
| amplitude height | 0.430 | -0.479 | 0.318 | 0.172 |
| amplitude width | 0.085 | -0.281 | -0.655 | 0.676 |
| amplitude symmetry | -0.448 | -0.529 | -0.065 | -0.106 |
| time to peak | 0.466 | 0.505 | -0.104 | 0.185 |
| tCNR | 0.551 | -0.379 | 0.228 | -0.119 |

Loadings above 0.5 for each PC are highlighted in grey. Only PC1 and PC2 contribute to the separation between 7T and 3T measurements (indicated by the red frame).

### Supplementary results

#### Identification of RSNs

Using various similarity measures to two different sets of templates (Smith and Stanford, see methods) we reliably could identify nine RSNs (Extended Data Fig. 4a). Using the criteria of maximum similarity converging in both directions (template to ICA and ICA to template), five RSNs could be unambiguously assigned to both, the Smith Template and the Stanford template. These RSNs were the auditory (auN), primary visual (pVN), occipital visual (oVN), left and right executive control (LECN, RECN, respectively) networks. Sensorimotor (SMN) and anterior Saliency (aSN) were identified solely according to the Stanford template. The default mode network (DMN) was assigned with high similarity to the corresponding Smith template, but the best match of the Stanford template to this ICA component by the similarity measures was the precuneus network. The precuneus network and the DMN strongly overlap and may not be separated accordingly using 20 ICA components as used for both the Smith template and our data. However, the Stanford template was created using 30 ICA components, therefore the DMN was separated into a ventral and a dorsal part and additionally the precuneus network was detected. The similarity to the templates varied greatly between RSNs, indicating RSN specific variations in spatial reliability. Highest spatial correlation to the Smith template was found for the 3T pVN (fisher's $z = 0.896$), the least similarity occurred for the 7T AuN (fisher's $z = 0.234$) with a median of fisher's $z=0.484$ or all identified RSNs (Extended Data Fig. 4b). The spatial overlap of the binarized RSNs to the Stanford template, i.e. the proportion of template ROIs covered by the RSNs, was also highest for the 3T pVN (overlap ratio = 0.997) and lowest for the 7T lVN (overlap ratio = 0.181) (Extended Data Fig. 4c). The lVN of the Stanford template covers dominantly regions involved in visuospatial processing, whereas the lVN of the Smith template is located lateral occipital and associated more to the higher visual areas. The median overlap was nearly 60 % (overlap ratio = 0.595). According to often greater extension of the binarized 3T RSNs most overlap values were also higher for 3T. The similarity according to the Jaccard index, which normalizes for the RSN size, showed no difference for the majority of RSNs between 3T and 7T (Extended Data Fig. 4d). The same could be observed regarding the spatial correlations to the Smith templates (Extended Data Fig. 5b). Thus, variations in similarity to the template were not dependent on field strength but rather on the intrinsic variability of the RSNs themselves.

**Quality assessment of RS graphs**

Quality assessment was initially demonstrated by the dominant connectivity strength between bilateral regions, visible as diagonal lines in the correlation matrices with brain regions sorted by hemisphere (Extended Data Fig. 5a). Mathematically, the data quality was assessed by the linear fit of the percentage correlation rank of corresponding bilateral brain regions in dependence on their anatomical distance (Extended Fig. 5b). Though the percentage rank difference per cm anatomical distance was near 0 (-0.25% for 3T and -1.3 % for 7T), the distance dependency was significantly higher for 7T data (Extended Data Fig. 5b,c, paired t-test, $p<0.05$). However, no field strength specific significant differences in the average correlation rank of all corresponding bilateral brain regions could be observed indicating comparable basic data quality for both field strength (Extended Data Fig. 5d).

### Supplementary discussion

**Early memory consolidation is influenced by specific anticorrelated connectivity modulations**

Another important feature of memory consolidation is the switch between task positive and task negative RS networks, giving anticorrelated connections a functional meaning[44-47]. Here, we found modulations of anticorrelated connections only for MFG, but not for M1. The MFG is the core region of the task positive executive control network which is also implicated in working memory processing[72,73], thereby acting as the main antagonist to the task negative DMN[74]. We found both, enhanced and reduced anticorrelated functional connectivity strength. This is in line with Piccoli et al. (2015)[47], who suggest a dynamic switch of FC between task positive and task negative (i.e. the DMN) brain networks during memory consolidation. In general, enhanced anticorrelation is associated with better memory performance[44-46]. In this context we found enhanced anticorrelation to MFG in contralateral regions specifically important for memory consolidation (angular gyrus[75,76], inferior temporal gyrus[77,78], occipital fusiform gyrus[79,80] and parahippocampal gyrus[81,82]), as well as the ipsilateral superior posterior cerebellum, namely lobule VI, Crus II, Crus I and lobule VIIb. The cerebellum is not only involved in motor control but also in cognitive and emotional functions including working memory[83-86]. Recent human studies revealed a functional topography of the cerebellum with distinct representation of different sensorimotor and cognitive functions[84,87]. Some of these functional compartments overlap, such as language and emotion[84], motor and sensory, as well as language and working memory[87], indicating an integrative function of these overlapping areas. However, none of these studies investigated the overlap of motor response and working memory, although the description of the activated areas for both tasks separately indicate the possibility of such an overlap. Consistently, finger movement activated the ipsilateral anterior lobule V and VI, while activation of areas involved in working memory were located in the right lobule VI and, depending on the memory load, extended to right CrusII and VIIb[84,87,88]. In the present work, we found enhanced anticorrelation to the memory related MFG for exactly those cerebellar regions (right lobule VI, CrusII and VIIb) that count for working memory. However, additionally to the described motor related lobule V and VI these regions were also activated during task performance. Presuming a spatial overlap of cerebellar motor control and memory function support the hypothesis that the replay of task related activity during rest is directly coupled to working memory circuits highlighting the integrative role of the RS itself in continuous integration of behavior and memory.

Reduced anticorrelations are more difficult to interpret. However, it stands out, that those structures that show reduced anticorrelation on the left hemisphere were strengthened in their positive correlation on the ipsilateral side (Au, S2, MTE) and vice versa (ITE). Maybe this interhemispheric interplay promotes the lateralized region's specific function in memory consolidation. This functional memory related lateralization has been frequently observed with memory task induced activation patterns[36] and recently also for anticorrelated connectivity strength[46].

**Early memory consolidation is influenced by specific hemispherical lateralization**

Specifically for episodic memory and the prefrontal cortex a concept of a hemispheric encoding/retrieval asymmetry (HERA) has risen decades ago[49,51]. This model describes a stronger activation of the left prefrontal cortex during encoding of information into memory and the right prefrontal cortex being more active during retrieval of memory information. This could be recently verified for verbal memory tasks, but not for visual[48] and was also observed in rats performing a novel object recognition test[50]. In the present study, a HERA like switch between hemispheres was observed for M1, which was activated on the left hemisphere during finger-tapping (respective encoding) and showed enhanced connectivity strength to memory related regions on the right hemisphere in subsequent rest (presumed to be an early maintenance phase of memory consolidation). Additionally, the right MFG, which is part of the prefrontal cortex, showed significantly modulated connectivity including anticorrelation in an early maintenance phase of memory consolidation, although it was activated bilaterally during encoding. In light of the present findings the following question regarding the functional lateralization during memory consolidation arise:

1. Does the hemisphere switch observed on memory retrieval already takes place in the early maintenance phase directly after encoding?

2. Is the hemispheric switch itself actually a fundamental property of memory consolidation or does the activation during memory encoding not primarily depend on the asymmetry of the brain function when coping with specific tasks? It should be noted here that the HERA principle is observed primarily in verbal tasks and not in visual ones. Verbal processing in the brain is intrinsically asymmetrical, while visual tasks can generally be assumed to be bilateral. The unilateral motor task performed here, on the other hand like the verbal tasks, also leads to an asymmetrical activation and a hemisphere switch.

3. Does a possible hemisphere switch between memory encoding and early maintenance apply to all regions that are dominantly involved in asymmetrical processing of sensorimotor or cognitive experiences? Here, we were able to demonstrate this for the motor cortex in context of a motor task; the possible transfer of this principle for example to unilateral auditory tasks involving the auditory cortex would be interesting.
